# A Novel Actinobacteria-Specific Ribosome Hibernation Factor in *Mycobacterium tuberculosis*

**DOI:** 10.1101/2024.11.25.625233

**Authors:** Soneya Majumdar, Yunlong Li, Swati R. Manjari, Howard B. Gamper, Ya-Ming Hou, Nilesh K. Banavali, Anil K. Ojha, Rajendra K. Agrawal

## Abstract

*Mycobacterium tuberculosis* (Mtb), the causative agent of tuberculosis (TB), persists within its host for long periods. Treatment of TB requires prolonged administration of antibiotics, some of which target Mtb ribosomes, the protein synthesizing machine of the cell. We determined high-resolution cryo-EM structures of Mtb ribosomes obtained from biofilms under zinc-depleted conditions that are expected to induce ribosome hibernation. In these structures, in addition to finding the Mtb homolog of the hibernation promoting factor Mpy that binds to the decoding center in the small ribosomal subunit, we discovered an actinobacteria-specific protein, previously of unknown function, bound in the nascent polypeptide-exit tunnel (NPET) within the large ribosomal subunit. We refer to this NPET-bound protein as a actinobacteria-specific ribosome tunnel <u>o</u>cclusion factor (RTOF). RTOF is found with vacant or E-tRNA-bound 70S ribosomes, but not in actively translating Mtb ribosomes, suggesting that its binding coincides with ribosome hibernation. RTOF makes strong interactions through the entire stretch of NPET extending up to the peptidyl-transferase center (PTC), such that it would block the passage of nascent polypeptide chain. The highly conserved FRRKSG motif of RTOF also interferes with accommodation of the formyl-methionyl moiety of initiator fMet-tRNA^fMet^ in the conformation required for translation initiation. A molar excess of mRNA and initiator fMet-tRNA^fMet^ displaces RTOF by triggering a significant alteration to the conformation of FRRKSG motif, which suggests how RTOF-bound hibernating ribosomes can be reactivated under optimal cellular conditions. The presence of RTOF would interfere with the binding of PTC-targeting drugs, such as macrolides, lincosamides, and oxazolidines, thereby maintaining a drug-free pool of hibernating Mtb ribosomes.

## Introduction

Tuberculosis (TB), caused by the *Mycobacterium tuberculosis* (Mtb), remains one of the world’s leading causes of death ^1^. Treating TB requires daily administration of multiple antibiotics for six months^2^. This prolonged treatment is partly attributed to a persister subpopulation of Mtb cells exhibiting phenotypic drug tolerance, presumably due to their non-replicating and slow metabolic state ^3–6^. The molecular mechanisms resulting in non-replicating persisters of Mtb in hosts remain unclear. It can be reasonably assumed that Mtb persisters develop under conditions that limit the growth of Mtb in lung microenvironments, such as hypoxia, metal deficiency, or host-induced toxicity^3^. For example, limiting concentrations of free zinc, likely prevalent in the sputum and lung lesions of Mtb-infected humans^7^, induces hibernation of ribosomes in Mtb^8^, implying that persistence through ribosome hibernation is a likely scenario *in vivo*.

Ribosomes are large multicomponent ribonucleoprotein complexes that catalyze protein synthesis. Since protein synthesis consumes over 40% of cellular energy^9,10^, growth inhibitory conditions trigger cellular responses to down-regulate protein synthesis and ribosome biogenesis. For growth recovery upon nutrient replenishment, non-replicating cells must maintain a pool of intact, functionally competent, non-translating ribosomes that can quickly resume protein synthesis and kick-start the metabolic pathways^11^. This is achieved through ribosome hibernation, which involves the reversible inactivation and stabilization of ribosomes primarily through the binding of specialized proteins that prevent binding of canonical ligands such as mRNA and tRNAs ^12–14^. Genetic knockouts of hibernation factors accumulate defective ribosomes that fail to reactivate when favorable conditions return ^15–17^. The binding of hibernation factors to the ribosome prevents translation and shields the active centers of ribosome from cellular nucleases or proteases^12–17^.

Ribosome hibernation is a conserved process in both bacteria and eukaryotes, with several hibernation factors reported in eukaryotes^18–22^ and a few in bacteria^23–26^. In bacteria, three major families of hibernation factors have been identified: (i) the pY family protein (including long-HPF, short-HPF, and YfiA) which occupies the mRNA binding channel and the A and P sites on the small (30S) ribosomal subunit (SSU)^23–25^; (ii) RMF, found only in γ-proteobacteria, partners with short-HPF to form hibernating 100S ribosomes and binds at the binding site of the 5’-end of the mRNA ^26^; and (iii) a recently proposed hibernation factor called Balon, which binds near the A site of 70S ribosomes ^27^.

*Mtb* contains two homologs of the pY family of proteins, Rv0079 and Rv3241c ^11^. Rv0079, induced in response to hypoxia, stabilizes the SSU and is essential for preserving cellular viability in a growth-arrested state. It has been shown that Rv3241c and its *M. smegmatis* (Msm) ortholog, MSMEG_1878, hibernate ribosomes under growth-limiting condition produced by limiting concentrations of zinc^8,28^. We reported the first high-resolution structure of a hibernating *M. smegmatis* ribosome, revealing that, unlike other long-HPFs, Mpy hibernates ribosomes in a 70S state ^8^. Mpy binds at the decoding region via its N-terminal domain, preventing the binding of mRNA and tRNAs. Our structure also revealed that Mpy prevents conformational changes in the 16S rRNA helix 44 (h44) that is needed to facilitate the binding of aminoglycoside antibiotics. Thus, in addition to stabilizing translationally stalled ribosomes, Mpy also protects ribosomes from aminoglycoside antibiotics^8^. Another unique mode of ribosome hibernation was reported in mycobacteria involving Rv2629 and MSMEG_1130 as the Balon orthologs in Mtb and Msm, respectively^27^. Balon can bind near the A site of both vacant and actively translating ribosomes and is recruited by elongation factor-Tu (EF-Tu) in the GDP-bound state. The proposed mechanism of Balon-mediated ribosome hibernation that presumably targets a stalled translation-elongation complex contrasts sharply with the other known mechanisms, wherein the substrates for hibernation factors are typically non-translating ribosomes.

The studies mentioned above suggest that ribosome hibernation in mycobacteria can be achieved through diverse mechanisms, but the atomic structure of hibernating ribosomes in Mtb remains unresolved. Here, we report cryo-EM structures of hibernating ribosomes from biofilm cultures of Mtb, which reveal a new actinomycete-specific protein that binds at the nascent peptide exit tunnel (NPET). This protein, referred to as the ribosome exit tunnel occlusion factor (RTOF), occupies vacant 70S or E-tRNA-bound 70S ribosomes, but not translation elongation complexes with A- and P-site tRNAs. RTOF also associates with the NPET of the dissociated large (50S) ribosomal subunits (LSUs). Similar NPET-associated hibernation factors have been reported in eukaryotes, but this is the first occurrence found in bacteria, which reveals a new mechanism of bacterial ribosome hibernation.

## Results

### Identification of a novel hibernation factor in Mtb biofilm ribosomes

We isolated ribosomes from a 7-week Mtb biofilm culture grown under zinc-depleted (C-) conditions ^8,28^. This growth condition has been previously found to induce ribosome hibernation by Mpy^8^. Consistent with our previous studies in Msm, we observed that Mtb ribosomes under these conditions were assembled with the paralogs of S14, S18, L28, and L33 that lack the zinc-binding motif CXXC (**fig. S1A, B**). In contrast to Msm, Mtb contains two C- paralogs of the ribosomal protein L28, Rv2058c and Rv0105c (**fig. S1C**). Our structures reveal a distinct density corresponding to the Rv2058c paralog, which is consistent with its presence in the same operon as the other C- paralogs: Rv2055c (S18), Rv2056c (S14), and Rv2057c (L33).

Cryo-EM analysis revealed that only ∼14% of the 70S ribosome population from the biofilm was in an actively translating state, i. e. bound to either P- or both A- and P-site tRNAs (**Figure 1A**). The remaining 86% of the ribosomes were either vacant (**fig. S2A**), carried only an E-tRNA (**fig. S2B**), or were occupied by Mpy (**Figure 1B**). We observed a density corresponding to a small protein inserted into the NPET of the 72% of the non-translating 70S ribosome population (**Figure 1A**). Further classification of the 70S population containing this density revealed three distinct sub-classes (**fig. S3**): (i) a vacant 70S complex (**fig. S2A**), (ii) a 70S-E-tRNA complex (**fig. S2B**), and (iii) a 70S-Mpy complex (**Figure 1B, fig. S2D, E**). Each of these classes were refined to resolutions of 2.8Å, 2.7Å, and 2.7Å, respectively (**fig. S3, S4**).

**Figure 1.**
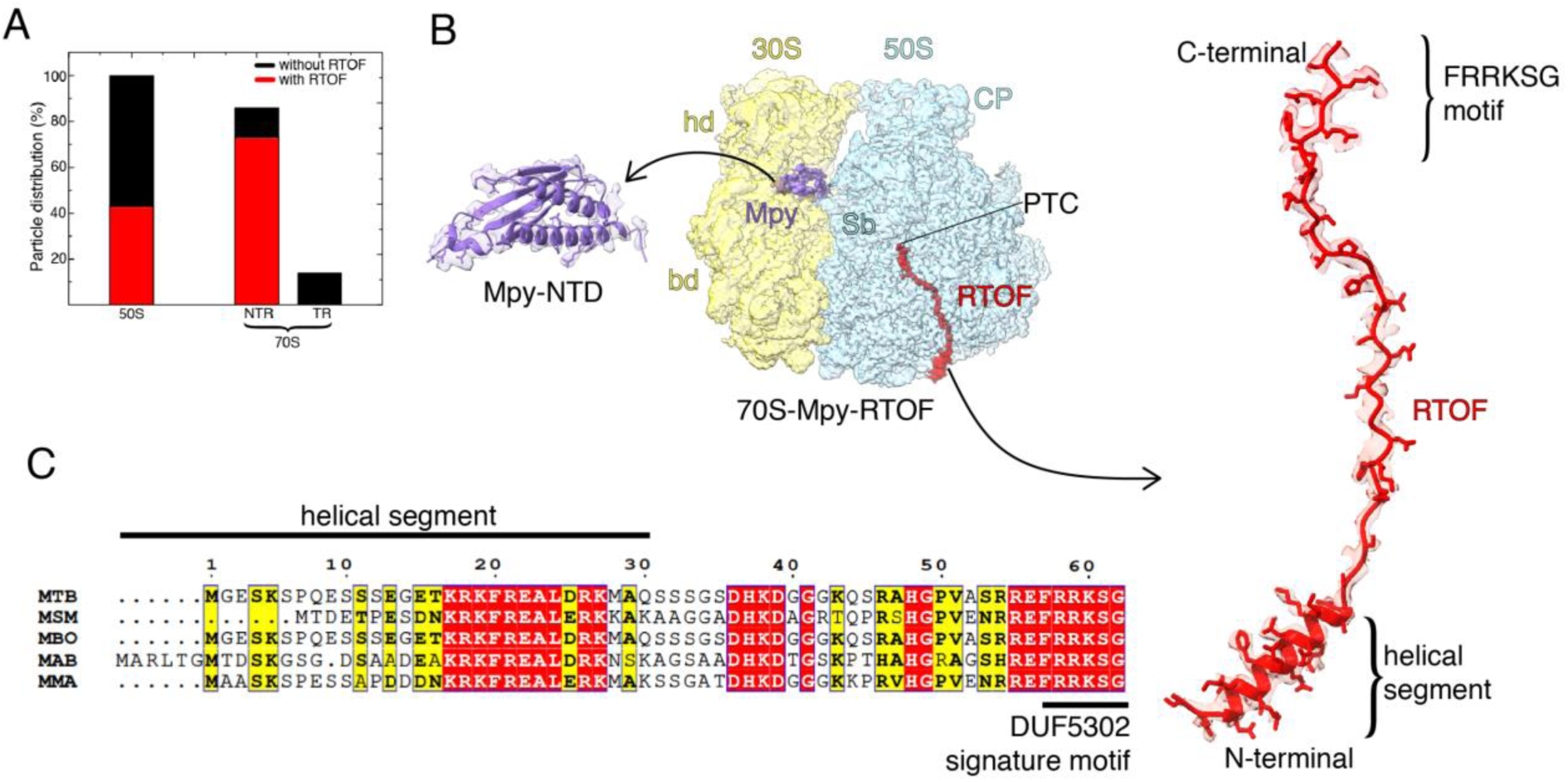
RTOF is a hibernation factor that associates with non-translating 70S ribosomal subunits. (A) Plot showing the percent distribution of RTOF-bound (red) and unbound (black), 50S, non-translating 70S (NTR), and translating 70S (TR) ribosomal subunits. (B) A 2.7 Å cryo-EM structure of the 70S-Mpy-RTOF complex, highlighting well-defined densities for RTOF (red) and Mpy (violet). Structural landmarks for the 30S small subunit (semitransparent yellow): hd, head; bd, body; and for the 50S large subunit (semitransparent blue): CP, central protuberance; PTC, location of peptidyl transferase center; and Sb, L7/L12 stalk base. (C) Sequence alignment of RTOF homologs from *Mycobacterium tuberculosis* (MTB), *Mycobacterium smegmatis* (MSM), *Mycobacterium bovis* (MBO), *Mycobacterium abscessus* (MAB), and *Mycobacterium marinum* (MMA). Identical amino-acid residues are highlighted in red, and similar residues are highlighted in yellow. The N-terminus helical domain and DUF5302 signature motif at the C-terminus are indicated.

To identify this NPET-inserted protein density, we first fit a poly-alanine model into it. This model appeared to be approximately 60 amino acids long, featuring a small N-terminal alpha helix emerging from the NPET and a long unstructured region extending up to the peptidyl-transferase center (PTC) within the LSU (**Figure 1B**). We then examined proteins smaller than 8 kDa in the mass spectrometry data obtained from the same ribosome sample for an independent study (Corro et. al., manuscript communicated). Among these, we identified an uncharacterized protein of 6.8 kDa that appeared to be a promising candidate. The Alphafold structure ^29,30^ of this candidate protein closely resembled our poly-alanine model having a short N-terminal alpha-helix followed by a longer unstructured region. The amino acid sidechains in the sequence also matched perfectly with the sidechain densities in the map. This confirmed that the protein, which we named the ribosome tunnel occlusion factor (RTOF) (**Figure 1B**), was encoded by the gene Rv1155a.

The three RTOF-bound ribosome structures show that RTOF preferentially associates with the NPET of non-translating 70S and hibernating Mpy-bound 70S monosomes (**Figure 1A-B**, **fig S2**). For the third structure, we confirmed that the bound Mpy is indeed encoded by Rv3241c, ruling out the presence of its close homolog Rv0079 **(fig. S2D, E)**. This is consistent with our previous study in Msm^8^, which demonstrated that Mpy is recruited onto the ribosome under zinc-depleted conditions. We also found RTOF in complex with 43% of the free Mtb LSUs (**fig. S2C**). Ribosomes isolated from *Mtb* biofilms thus carry an additional protein that exhibits two key characteristics of a hibernation factor: it associates with a major population of non-translating ribosomes under stress conditions, and occupies an active site of the ribosome.

### RTOF homologs are specific to Actinobacteria

RTOF is a previously uncharacterized protein, encoded by the Rv1155a gene, belonging to the <u>D</u>omain of <u>U</u>nknown <u>F</u>unction 5302 (DUF5302) protein family. Homologs of RTOF are found across mycobacterial species. Alignment of RTOF sequences in mycobacteria reveals high (63%) sequence similarity (**Figure 1C**) characterized by a highly conserved FRRKSG motif at its C-terminal end (**Figure 1C**), which suggests a crucial functional role for this sequence. Interestingly, the protein length varies through extensions at their N-termini (**Figure 1C)**, which emerges from the sovent side of NPET. The InterPro database (InterPro entry: IPR035172) shows that RTOF is specific to the Actinobacteria **(fig. S5)**. Most species, including the mycobacterial species, contain one copy of this protein while some Streptomyces and Actinokineospora species encode two copies.

RTOF is rich in charged amino acids (39%) which allows it to form multiple electrostatic interactions with the 23S rRNA and ribosomal proteins, uL22, uL23, uL29, that contribute to formation of the NPET (**Figure 2A**). The binding orientation of RTOF within the tunnel mirrors that of a nascent peptide, i. e. it is bound with its initially synthesized N-terminus at the NPET exit site and its later synthesized C-terminal end at the PTC. This suggests that it can block the NPET immediately upon its synthesis by the ribosome rather than first getting post-translationally released and then re-binding to the ribosome (**Figure 2A**). We observe a significant population of LSUs bound to RTOF (**fig. S2 C**), which could be a result of 70S-RTOF recycling by the ribosome recycling factor (RRF) and elongation factor-G (EF-G)^14,31^. The possibility that RTOF could also bind post-translationally to the 70S ribosomes and LSUs leading to ribosome hibernation cannot, however, be excluded.

**Figure 2.**
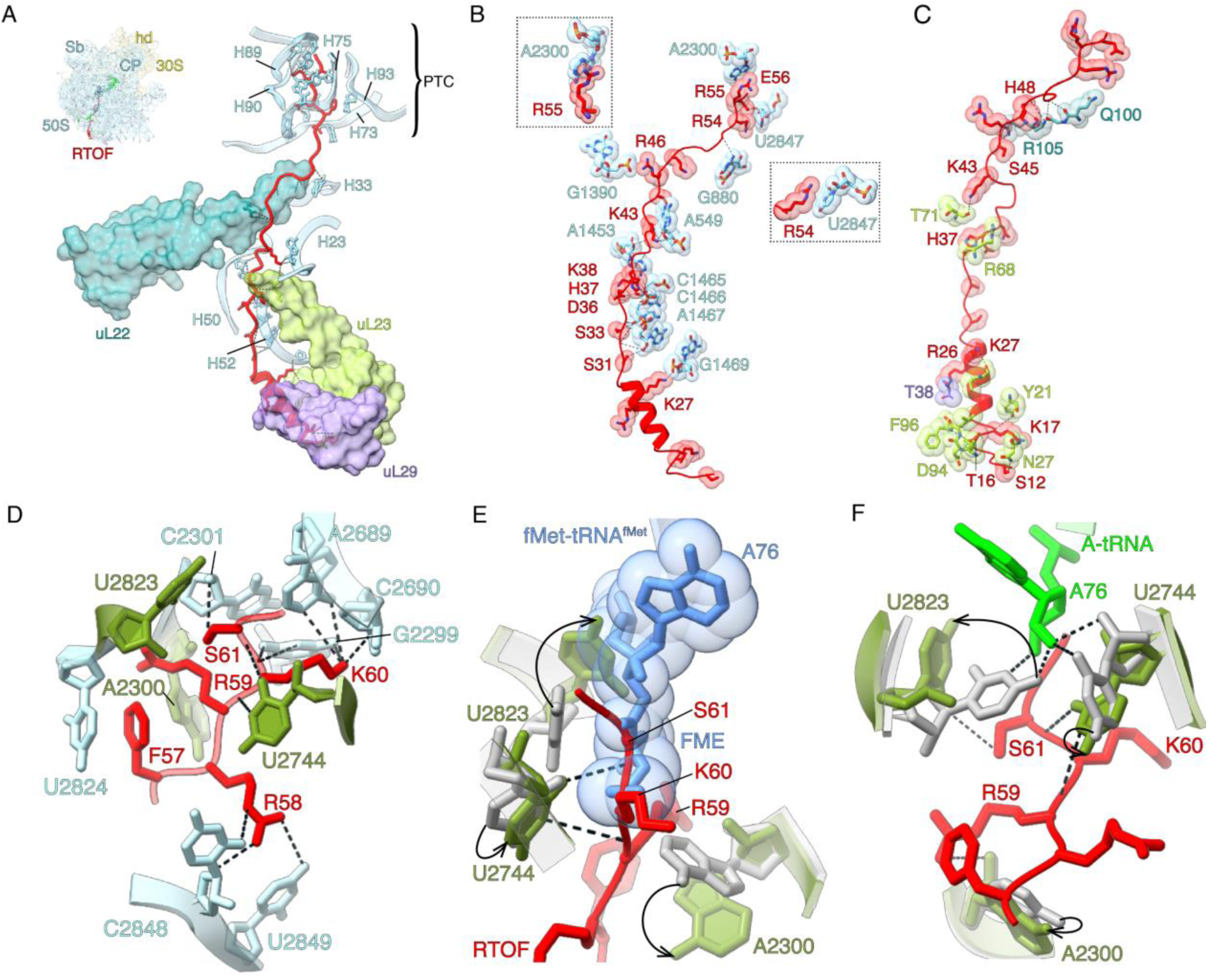
RTOF interacts with components of the NPET and the PTC, inducing conformational changes in some of the key PTC residues. (A) Components of the NPET, including ribosomal proteins uL22 (teal), uL23 (yellow-green), uL29 (purple), and multiple rRNA helices identified by helix (H) numbers (cyan) that interact with RTOF (red), are displayed. (B) Specific interactions between RTOF amino acids and 23S rRNA nucleotides are shown. Two boxed zoomed-in views highlight stacking interactions between R55 and A2300, and between R54 and U2847. (C) Interactions between RTOF amino acids and ribosomal proteins are shown. The r-protein color scheme is consistent with panel A. (D) Interactions of the DUF5302 signature motif, FRRKSG (red), with the PTC components. Three PTC nucleotides—A2300, U2744, and U2823 (olive green)—undergo significant conformational changes to accommodate RTOF. (E) Superimposition of the 70S-RTOF structure with the 70S-fMet-tRNA^fMet^ complex (PDB: 7MT2, ^57^). Adenine residue (A76) and fMet moiety of fMet-tRNA^fMet^ (cornflower blue) and associated bases of the 23S rRNA, A2300, U2744, and U2823 (grey), indicate their positions in a 70S initiation complex. Conformational changes in these PTC bases are illustrated with curved arrows. (F) Superimposition of the 70S-RTOF structure with the *E. coli* 70S-A-P-tRNA complex (PDB: 7K00, ^36^). Adenine residue of A-tRNA (lime green) and its interactions with U2744 and U2823 (grey) are shown. The conformational changes in the three PTC bases upon RTOF binding (olive green) are again depicted with curved arrows.

### RTOF locks the PTC in an inactive conformation

RTOF is absent from the 14% of the 70S ribosome population that carried either a P- or both A- and P-site tRNAs, and mRNA, indicating that it does not bind to an actively translating ribosome. To understand how RTOF stably binds to and occludes the NPET, we analyzed its molecular interactions within the tunnel. RTOF interacts extensively along the entire length of the NPET (**Figure 2B, C**). RTOF, being rich in charged amino acids, forms numerous electrostatic interactions with the 23S rRNA (**Figure 2B**) and the r-proteins lining the lower 2/3^rd^ of NPET (**Figure 2C**). Specifically, RTOF residues K27, S31-33, H37, D36, K43, R46, and E56 interact with the 23S rRNA nucleotides G1469, A1467, C1466, C1465, A1453, A549, G1390, and U2847, respectively (**Figure 2B**). The RTOF residues S45, H48 interact with Q100 and R105, respectively, of r-protein uL22; K43, H37, K27, K17, T16 and S12 interact with T71, R68, Y21, D94, F96 and N27, respectively, of r-protein uL23; and R26 with T38 of r-protein uL29 (**Figure 2C**). Several hydrogen bonds also form between the backbone of RTOF and the NPET components. Further, two RTOF arginine residues, R54 and R55, engage in π-stacking interactions with A2300 and U2847 of the 23S rRNA (**Figure 2C, zoomed in view**).

Ribosome hibernation factors bind to ribosomes in response to stress and inhibit protein synthesis^32^. We analyzed the 70S-RTOF structures to understand how binding of RTOF could inhibit protein synthesis. The extensive interactions of the highly conserved C- terminal FRRKSG motif of RTOF with the PTC could deter translation initiation (**Figure 2D**). This motif interacts with several crucial PTC residues, and its accommodation in the PTC is accompanied by an induced fit that involves conformational changes in three PTC residues^33,34^: A2300 (A2062 in *E. coli*), U2744 (U2506 in *E. coli*), and U2823 (U2585 in *E. coli*) (**Figure 2D**). U2823 and U2744 are essential for the proper positioning of the CCA ends of the initiator tRNA and the A-site tRNA during the first peptide-bond formation at the PTC^33,35^.

To understand how these interactions might affect the accommodation of A- and P-tRNAs, we superimposed the structures of 70S-E-tRNA-RTOF and 70S-A-P-tRNA^36^ (**Figure 2E, F**). We found that there is no direct steric overlap between the A-tRNA and RTOF. RTOF residues K60 and S61 do occupy the same location as the formyl-methionyl (fMet) moiety of fMet-tRNA^fMet^. The residue S61 of the FRRKSG motif also interacts with U2823, holding it in a conformation that would overlap with A76 (of CCA) of the P-tRNA (**Figure 2E**). Both S61 and R59 interact with U2744, stabilizing it in a conformation that would prevent interaction with the CCA end of A-tRNA (**Figure 2F**). These analyses suggests that RTOF could act as a competitive inhibitor of translation, likely affecting the accommodation of fMet-tRNA^fMet^ at the translation initiation step.

### Comparison of RTOF with PTC-associated inhibitors of translation

Several proline-rich antimicrobial peptides, synthesized as part of the innate immune response in plants, insects, and animals, bind to the NPET of the ribosome, inhibiting protein synthesis ^37,38^. These peptides disrupt various stages of protein synthesis depending on their interaction with the PTC. For instance, Bactenecin-7 (Bac7) binds to the NPET in a reverse orientation as compared to a nascent polypeptide. A superimposition of the *Thermus thermophilus* 70S- Bac7(1-16) and 70S-RTOF structures (**Figure 3A, B**) reveals that while RTOF overlaps with the fMet moiety of fMet-tRNA^fMet^, Bac7 (1-16) overlaps with the binding site of the CCA-end of A-tRNA (**Figure 3A, B**). Biochemical studies have shown that Bac7 primarily inhibits translation by blocking the accommodation of aminoacyl-tRNA at the A site during translation elongation, although at higher concentrations, it can also affect translation initiation ^38^. Drawing parallels with these findings, we conclude that RTOF likely inhibits protein synthesis by blocking translation initiation.

**Figure 3.**
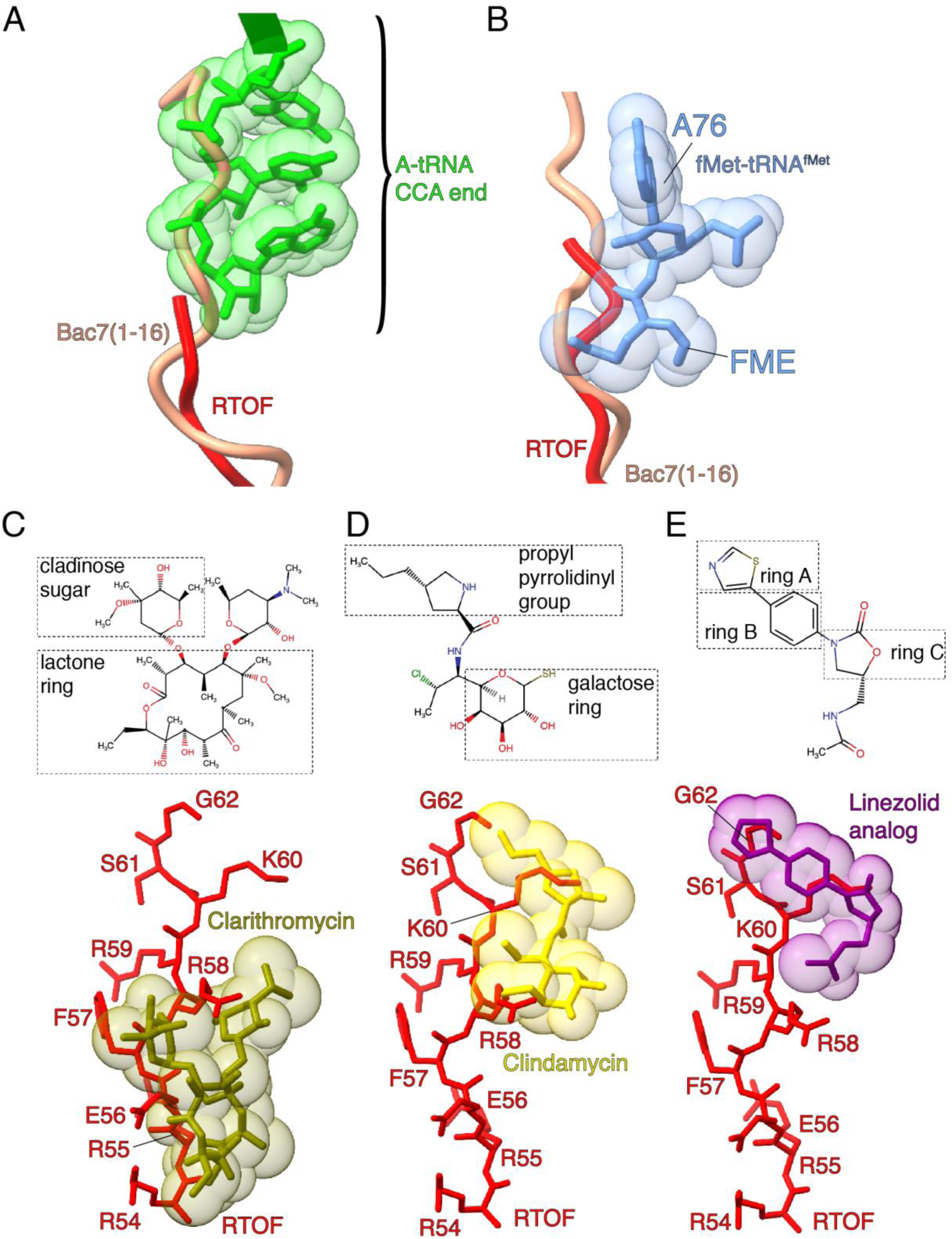
RTOF inhibits translation initiation and impacts the binding of PTC-targeting antibiotics. (A) Superimposition of the 70S-RTOF structure with the *E. coli* 70S-A-tRNA (PDB: 7K00, ^36^) complex and *T. thermophilus* 70S-Bac7 (PDB: 5F8K, ^38^) structures. The CCA end of A-tRNA (lime green) overlaps with Bac7 (salmon) but not with RTOF (red). (B) Superimposition of the 70S-RTOF structure with the Mtb 70S-fMet-tRNA^fMet^ complex (PDB: 7MT2, ^57^) and *T. thermophilus* 70S-Bac7 structures (PDB: 5F8K). The formyl-methionyl group and A76 of the CCA end of fMet-tRNA^fMet^ (cornflower blue) show significant overlap with RTOF and a lesser overlap with Bac7. (C-D) Superimposition of the 70S-RTOF structure with the structures of (C) Mtb 50S-Clarithromycin (PDB:7F0D,^39^), (D) *E. coli* 50S- Clindamycin (PDB:8CGD, ^42^), and (E) 50S-Linezolid (PDB:5V7Q, ^40^). Different and/or overlapping segments of FRRKSG motif of RTOF show substantial overlap with clarithromycin (olive green), clindamycin (yellow), and linezolid (purple).

Clarithromycin, clindamycin, and linezolid, belonging to the macrolide, lincosamide, and oxazolidinone classes of drugs, respectively, are often used as second-line or adjunctive treatments for multi-drug resistant TB (MDR-TB) or infections caused by non-tuberculous mycobacteria (NTM). Given that RTOF extends across the NPET up to the PTC, we investigated whether it overlaps with the binding sites of these drugs. Superimposing the drug-bound 70S structures ^39–42^ with the 70S-RTOF structure revealed significant overlap between RTOF and bound drugs (**Figure 3C-E**). For clarithromycin, the C-terminal residues of RTOF (R54-R58) overlap with the cladinose sugar and part of the lactone ring (**Figure 3C**). In the case of clindamycin, the overlap occurs between RTOF residues R58-K60 and the galactose ring and pyrrolidinyl propyl group of the drug (**Figure 3D**). For linezolid, residues K60-G62 overlap with the oxazolidinone ring (ring A), as well as rings B and C (**Figure 3E**). Thus, the association of RTOF with the 70S or 50S ribosomes could potentially prevent binding, or affect the binding affinity, of these drugs.

### fMet-tRNA^fMet^ accommodation can displace RTOF from the NPET and reactivate protein synthesis

How hibernating ribosomes are reactivated when favorable conditions are restored is not fully understood. It is hypothesized that, during rapid growth, canonical protein synthesis ligands such as IF1 and IF3, which share overlapping binding sites with hibernation factors like HPF, may facilitate their displacement from the ribosome. Supporting this idea, *in vitro* assays have demonstrated that IF3 promotes HPF dissociation in *E. coli*^43^. Binding of RTOF to both non-translating 70S ribosomes and LSUs suggests that RTOF may remain associated with the LSUs even after the 70S ribosomes split into their individual subunits. We hypothesize that RTOF is either displaced upon the resumption of protein synthesis or actively removed through an alternative mechanism. To explore the first possibility, we assembled a complex of RTOF-bound ribosomes with fMet-tRNA^fMet^ and mRNA (Met-Phe) to evaluate how these ligands compete with RTOF for binding. Cryo-EM analyses of this complex revealed that 92% of the ribosomes contained P- and E-tRNAs, as compared to the 86% of vacant, E- tRNA-, or Mpy-bound ribosomes in control sample without the exogenous addition of mRNA and fMet-tRNA^fMet^ (**fig. S6, Figure 1A)**. Furthermore, we noted a significant reduction in RTOF-bound 70S particles from 73% to 31%, indicating that the addition of fMet-tRNA^fMet^ displaced RTOF from more than half of the ribosome population (**fig. S6, Figure 1A**).

To understand how the accommodation of fMet-tRNA^fMet^ leads to the displacement of RTOF, we refined the 70S-P-E-tRNA-RTOF and 70S-P-E-tRNAs bound structures, both to a resolution of 3.3Å (**Figure 4A, B, fig. S6, S7**). Although we observed a clear density for the fMet-tRNA^fMet^, including its CCA end in both structures, density for the fMet moiety in the RTOF-bound structure remained disordered (**Figure 4A, B**). Superimposition of the 70S- RTOF and 70S-P-E-tRNA-RTOF structures revealed that the FRRKSG motif of RTOF undergoes significant displacement in the presence of fMet-tRNA^fMet^ (**Figure 4C**). Specifically, F57 is displaced by 2Å, while R59 and S61 are shifted by 7.2Å and 3.6Å, respectively. The rest of the protein retains its conformation across the NPET (**Figure 4C**). The displaced position of the FRRKSG motif in the 70S-P-E-tRNA-RTOF structure possibly represents an intermediate state of fMet-tRNA^fMet^-mediated RTOF release from the ribosome.

**Figure 4.**
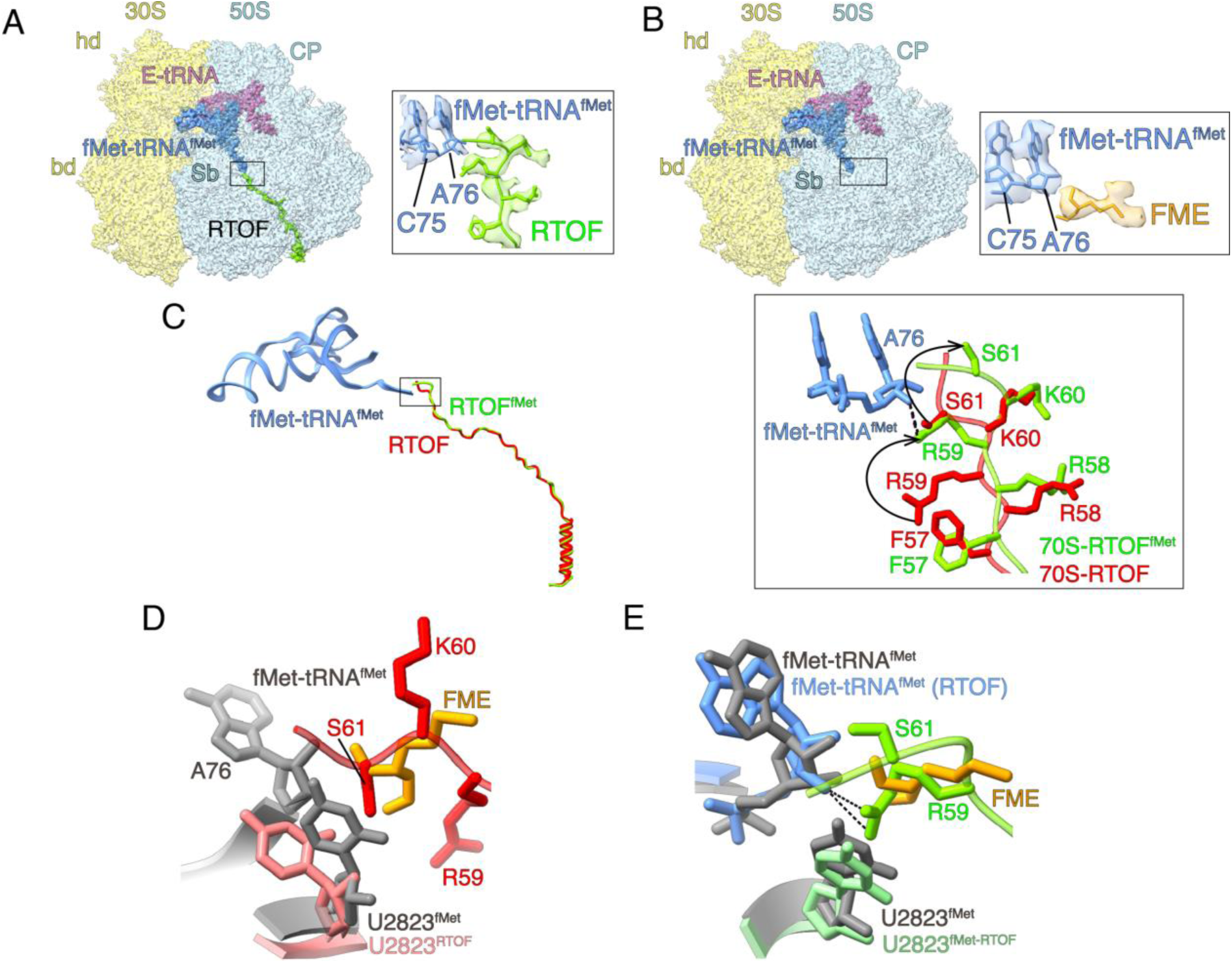
Accommodation of fMet-tRNA^fMet^ in the 70S-RTOF ribosome leads to partial displacement of RTOF. (A) A 3.3 Å cryo-EM structure of the 70S-fMet-tRNA^fMet^-RTOF complex, representing an intermediate state of RTOF (lime green) release from the ribosome. A boxed zoomed-in view highlights the co-accommodation of the fMet-tRNA^fMet^ CCA-end and the RTOF-FRRKSG motif, along with their respective densities. (B) A 3.3 Å cryo-EM structure of the 70S-fMet-tRNA^fMet^-E-tRNA complex. The zoomed-in view shows the densities corresponding to the CCA-end of fMet-tRNA^fMet^ and the formyl-methionyl group (FME, orange). (C) Superimposition of the 70S-RTOF and 70S-RTOF-fMet-tRNA^fMet^ complex structures, illustrating the conformational changes in RTOF (lime green) upon fMet-tRNA^fMet^ (cornflower blue) binding. The boxed zoomed-in view highlights the conformational changes, with curved arrows indicating significant alterations in RTOF residues, R59 and S61. (D) Superimposition of the 70S-RTOF and 70S-fMet-tRNA^fMet^-E- tRNA complex structures, showing the overlap of RTOF with the FME and the conformational shift of U2823 (from grey to salmon) between the RTOF-unbound and -bound states. (E) Superimposition of the 70S-RTOF and 70S-RTOF-fMet-tRNA^fMet^-E-tRNA complex structures, showing that despite the conformational changes in RTOF, it continues to overlap with the FME, U2823 reverts to its RTOF-unbound position (green to grey).

To understand how this altered conformation of the FRRKSG motif is co-accommodated with fMet-tRNA^fMet^, we compared the structures of the 70S-fMet-tRNA^fMet^, 70S-fMet-tRNA^fMet^-RTOF, and 70S-RTOF (**Figure 4D, E**). The comparison revealed that upon fMet-tRNA^fMet^ binding, the flipped conformation of the 23S rRNA residue U2823— induced by the interaction with RTOF S61, which would clash with the fMet-tRNA^fMet^ (**Figure 4D**)—is flipped back as S61 moves away (**Figure 4E**). Despite this, translation initiation would be inhibited in this intermediate state of RTOF because, even in the altered conformation, RTOF R59 of the FRRKSG motif interacts with the 3’-OH group of the ribose of A76 and occupies the position of the fMet moiety (**Figure 4E**). The absence of density for the fMet moiety when fMet-tRNA^fMet^ and RTOF are bound simultaneously to the ribosome (**Figure 4A**) suggest that the fMet moiety does not occupy a fixed position and hence is disordered in the map. Hence, the release of RTOF is essential for the formation of a functional post-initiation complex. These structural analyses suggest that (i) the presence of RTOF would deter translation at the initiation step, and (ii) binding of fMet-tRNA^fMet^ would initiate displacement of RTOF by forcing the FRRKSG motif to adopt an alternate conformation.

## Discussion

Bacteria enter a non-replicating state when faced with unfavorable environmental conditions, such as macro and micronutrient deprivation, antibiotic exposure, or osmotic stress ^44^. During this state, cells tightly regulate protein synthesis—an energy-intensive process ^10^—by storing ribosomes in an inactive but structurally and compositionally intact state. This phenomenon is considered to be crucial for cells to resume growth when favorable conditions return ^11^. Ribosome hibernation is a conserved process across both bacteria and eukaryotes. However, the structural and functional diversity among hibernation factors in bacteria makes it challenging to identify and predict the full spectrum of these factors. Previous studies in mycobacteria have shown that ribosomes hibernate in the 70S state (rather than the 100S state) upon binding of Rv0079 encoded RafH under hypoxia ^45^ and Rv3241c encoded Mpy under zinc-limiting conditions ^8,28^. Both these factors occupy the mRNA binding channel, spanning the A- and P-tRNA binding sites on SSU. In addition to the commonly accepted model that hibernation factors bind exclusively to non-translating ribosomes, recent findings suggest hibernation of actively translating 70S ribosome in mycobacteria by Rv2629-encoded Balon^27^, although the Balon function in other aspects of translation such as ribosome rescue remains to be fully elucidated. In this study, we identified a novel protein factor, RTOF, that binds to the NPET of the LSU and non-translating 70S ribosomes isolated from a 7-week Mtb biofilm culture grown under zinc-depleted conditions (**Figure 1B, fig. S1, S2)**.

Previous biochemical studies in *E. coli* have examined NPET-binding proteins like YqjD, ElaB, and YgaM ^46^. Unlike these proteins, which are approximately 100-110 amino acids in length, RTOF consists of only 62 amino acids. Despite sharing the NPET-binding function, RTOF exhibits poor sequence and structural similarity with YqjD/ElaB/YgaM (**fig. S8A)**. An InterPro database search indicates that while RTOF is unique to Actinobacteria, the YqjD/ElaB/YgaM family predominantly occurs in Proteobacteria, suggesting that RTOF and these proteins are functional analogs likely resulting from convergent evolution. RTOF also has key differences: (i) It binds as a monomer, whereas YqjD binds as a dimer; (ii) the orientation of RTOF within the NPET mimics a nascent polypeptide, while YqjD has an N-terminus interacting with the NPET and a C-terminus containing a transmembrane segment (**fig. S8B)**; (iii) RTOF extends throughout the NPET, from PTC to the exit of NPET, unlike YqjD, which only interacts with the tunnel exit. Importantly, while our structural analysis directly demonstrates that RTOF interacts with the NPET, the interaction of YqjD has been inferred from cross-linking studies rather than observed through molecular structures. These observations strongly support RTOF identification as a distinct and novel protein factor within Actinobacteria.

RTOF, discovered in this study, shows some similarities to eukaryotic NPET-binding hibernation factors such as Dapl1^22^ and MDF2 ^21^ (**fig. S9A**). In eukaryotes, these factors have been observed in two cases: (i) MDF2 in the metabolically inactive spores of microsporidia^21^ and (ii) Dap1b/Dapl1 in frog (Xenopus) oocytes^22^. These eukaryotic hibernation factors bind their ribosomes in coordination with additional factors—MDF2 associates with MDF1, which occupies the mRNA binding channel and E site, while Dap1b/Dapl1 binds along with eIF5a, eEF2, and Habp4, where Habp4 fills the mRNA channel, stabilizes eEF2 at the A site, and eIF5a occupies the space between the E and P sites of the ribosome (**fig. S9B)**. In contrast, ribosomes bound by RTOF in our study were either vacant, E-tRNA-bound, or Mpy-bound (**Figure 1B, fig. S1, S2**). Sequence analysis shows little conservation between RTOF, MDF2, and Dap1b, though all three are rich in charged amino acids (39%, 48%, and 27%, respectively), facilitating extensive interactions along the ribosomal NPET (**fig. S9A-C**). The orientation of these proteins within the NPET is consistent with co-translational binding mechanism upon their synthesis (**fig. S9D**). RTOF is significantly shorter—only 62 amino acids—compared to MDF2 and Dap1b, which are 160 and 113 amino acids, respectively, leading them to occupy different regions within the NPET and extending beyond it (**fig. S9A)**. RTOF extends precisely up to the position of the C-terminal residue of a typical nascent polypeptide chain, while Dap1b extends further by five amino acids into the ribosomal inter-subunit space (**fig. S9D**). MDF2, however, blocks the entire PTC, extending to occupy the P-tRNA acceptor arm (**fig. S9C, D**). Given that Dap1b and MDF2 are established hibernation factors that inhibit translation, the structural and functional similarities suggest a similar role for RTOF in Mtb.

The percentage of ribosomes carrying both Mpy and RTOF was relatively low (3%) as compared to 73% ribosomal population carrying exclusively RTOF. Moreover, all particles had C- r-protein paralogs, which is consistent with or previous findings that C+/C- ribosome remodeling precedes ribosome hibernation in zinc-depleted conditions^8,28^. Since these structures are derived from ribosomes isolated from an Mtb biofilm culture under zinc-depleted conditions without any externally added ligands, we believe that the observed classes represent *in vivo* states of the ribosome in this environment.

We also observed a significant percentage of LSUs bound to RTOF (**Figure 1A, fig. S2C**), likely resulting from the recycling or splitting of 70S-RTOF complexes by RRF-EFG^14,31^. This suggests that, in addition to its role as a hibernation factor, RTOF may also protect dissociated LSUs. Furthermore, when RTOF-bound ribosomes were incubated with Met-Phe mRNA and fMet-tRNA^fMet^, RTOF was displaced from a significant population (69%) of ribosomes without changing the overall population of RTOF-bound LSUs (compare **Figure 1A** and **fig. S6D**). The structure of the remaining population of the 70S-RTOF-fMet-tRNA^fMet^ complex revealed that the C-terminal FRRKSG motif of RTOF adopts an altered conformation in the presence of fMet-tRNA^fMet^, representing an intermediate state for RTOF release, (**Figure 4A, C**). The highly conserved FRRKSG motif (**Figure 1C**) likely serves as the primary anchoring point of RTOF on the ribosome. The observed conformational changes upon fMet-tRNA^fMet^ binding would compromise the anchoring interactions, thereby leading to the release of RTOF from the ribosome (**Figure 4C**).

Our findings allow us to propose the following model for RTOF-mediated 70S hibernation and LSU sequestration (**Figure 5**): In response to stress conditions, it is likely that RTOF binds to the NPET co-translationally (**Figure 5A→B**), although the possibility of post-translational binding of RTOF to the ribosome cannot be ruled out. The 70S-RTOF ribosomes may either be recycled by splitting factors into LSU-RTOF complexes and SSUs (**Figure 5C→F**) or remain intact as 70S-RTOF complexes (**Figure 5D**). These ribosome-RTOF complexes likely persist when the cell has depleted levels of fMet-tRNA^fMet^. Under growth-limiting conditions, including zinc starvation, Mpy also binds the ribosome to form a 70S- Mpy-RTOF complex (**Figure 5D→E**). Co-existence of RTOF and Mpy on the 70S ribosome will block all three major functional centers of the ribosome simultaneously: the decoding center, the PTC, and the NPET, effectively preventing futile translation cycles. However, when favorable conditions return and fMet-tRNA^fMet^ becomes available, Mpy would be recycled, probably with the help of RRF and EF-G (**Figure 5E→F**). The subsequent binding of canonical translation initiation factors and fMet-tRNA^fMet^ during translation initiation will displace RTOF (**Figure 5F→A**), allowing protein synthesis to resume.

**Figure 5.**
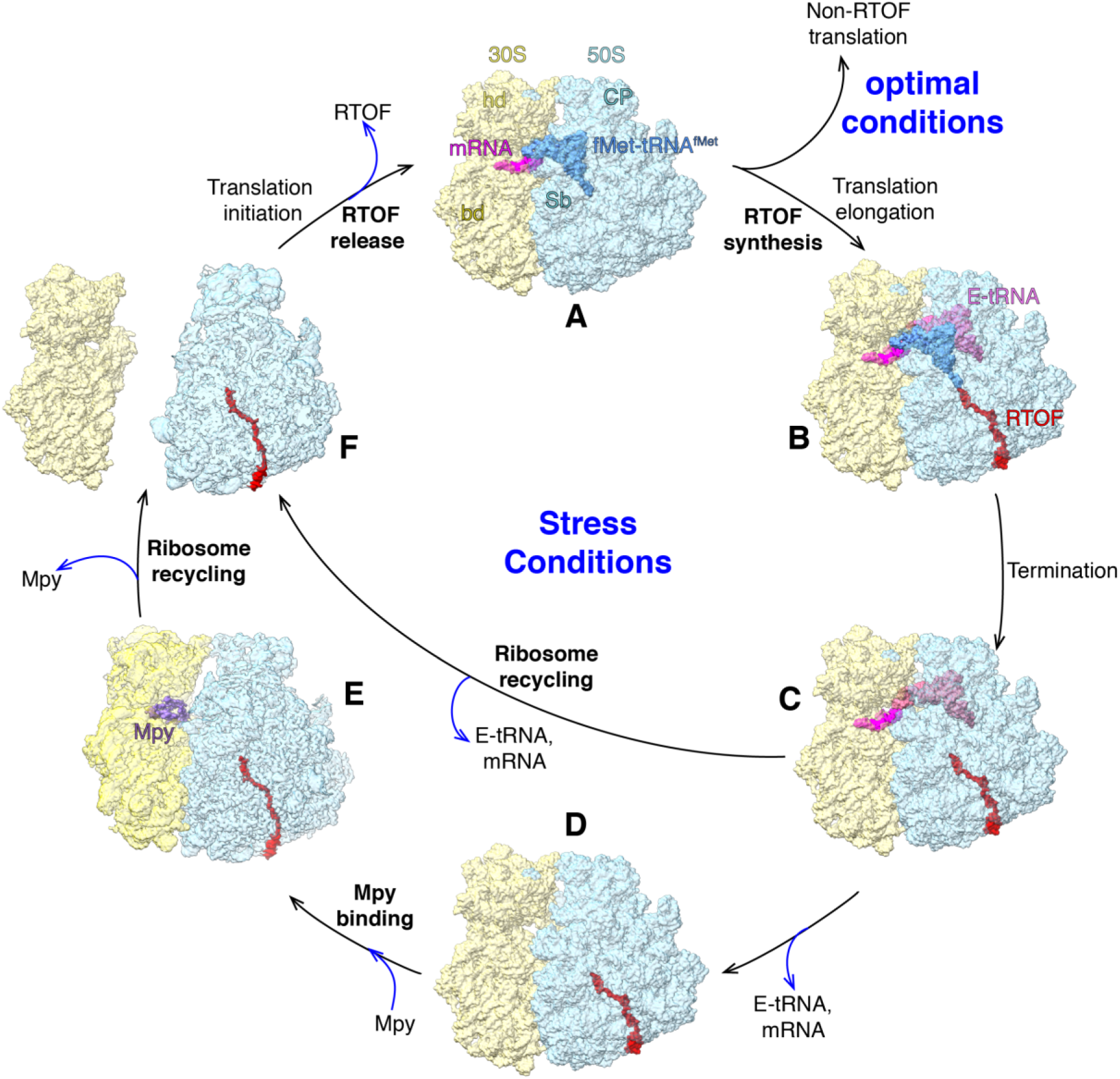
A proposed mechanism of co-translational RTOF recruitment to the Mtb ribosome during stress conditions. (A) Translation initiation complex, (B) translation of RTOF, (C) RTOF translation termination, leads to formation of mRNA-E-tRNA-RTOF post-termination complex (fig. S2B), (D) dissociation of E-tRNA and mRNA results in 70S-RTOF complexes (fig. S2A). (E) Binding of Mpy leads to formation of 70S-Mpy complex (Fig. 1B). (F) Recycling of 70S-Mpy complex would lead to removal of Mpy and dissociation of the 70S into 30S and RTOF-bound 50S subunits (fig. S2C). The panel (F) could also result from recycling of the post-termination complex shown in panel (C). RTOF would be eventually removed from 50S subunit during translation initiation complex formation (Fig. 4), which could either be on a stress-related mRNA in stress conditions, or on other non-stress related mRNAs under optimal growth conditions.

Macrolides, lincosamides, and oxazolidinones are classes of antibiotics that inhibit protein synthesis in bacteria. Macrolides such as azithromycin and clarithromycin bind to LSU, blocking the NPET and inhibiting protein synthesis^47^. Although macrolides are generally not first-line TB drugs due to intrinsic resistance mechanisms in Mtb, clarithromycin and azithromycin may be used in treatment regimens for non-tuberculous mycobacteria (NTM) infections like *Mycobacterium abscessus* (Mab)^48^, although induced resistance by these drugs remains a challenge. Although lincosamides such as clindamycin that bind near the aminoacyl moiety of the A-site tRNA of LSU are not very effective against mycobacteria due to intrinsic resistance^49,50^, linezolid, an oxazolidinone that binds to the 23S rRNA of LSU is effective against Mtb and is used as part of treatment regimens for multi-drug resistant (MDR) and extensively drug-resistant (XDR) TB^51^.

Given that a significant population of ribosomes from Mtb biofilms, which harbor drug tolerant persisters^52^, is bound to RTOF at the site that overlaps with those of binding sites of the macrolide, lincosamide, and oxazolidinone classes of drugs (**Figure 3C-E**), we propose that the presence of RTOF will help in drug-free sequestration of ribosomes in RTOF-mediated hibernating states. This suggests a new RTOF-mediated antibiotic resistance mechanism in pathogenic mycobacteria such as Mtb and likely in other mycobacteria including Mab.

## Methods

### Bacterial growth and ribosome purification

*M. tuberculosis* mc^2^7000^52^ was cultured as pellicles on air-medium interface in detergent-free Sauton’s media without supplemental zinc. After 7-weeks of growth, pellicles from four 12-well plates were spooled out in sterile 50mL conical tubes containing PBS with 0.05% (v/v) Tween-80 and centrifuged for 20 minutes at 8000 rpm and 4 °C in Thermo Scientific™ Fiberlite™ F12-6 × 500129. Cells were then flash frozen in liquid nitrogen until further use. Ribosomes were purified exactly as described previously^28^. Briefly, cells were pulverized frozen for in a mixer mill (Retsch MM400) for 3 minutes at 15 Hz for eight times. Twenty mL of ice-cold HMA-10 buffer (20 mM HEPES-K pH 7.5, 10 mM MgCl_2_, 30 mM NH_4_Cl, 5 mM β-mercaptoethanol) was added to the cell lysates and the mixture was centrifuged for 30 minutes at 4 °C in Thermo Scientific™ F21-8 x 50y fixed-angle rotor. The supernatant was treated with 3 units/mL RNase-free DNase (Ambion) for one hour at 4 °C. Crude ribosomes were separated from other low molecular weight cellular contents by centrifugation at 35000 rpm (126,000g) for 1 hour at 4 °C (Beckman rotor Type 70Ti). The crude ribosome pellet was resuspended in 10 mL HMA-10 buffer and layered on top of a 10 mL 32% sucrose solution in HMA-10 buffer and centrifuged for 16 hours at 37,000 rpm (140,000g) in a Beckman Type 70Ti rotor. The ribosome pellet was briefly rinsed and then solubilized in HMA-10 buffer and fractionated on 10-40% sucrose density gradient (SDG) as previously detailed to collect the 30S, 50S and 70S ribosomes. The ribosomes were quantified by measuring absorbance at 260 nm.

### Preparation of fMet-tRNA^fMet^

A culture of *E. coli* harboring a plasmid with an expression vector for fMet-tRNA, controlled by the LacI repressor, was grown until the optical density at 600 nm reached 0.4–0.6. At this point, IPTG was introduced to trigger tRNA expression. After 8 hours, the cells were collected, lysed using phenol, and total tRNA was isolated through a series of differential precipitation steps ^53^. Roughly half of the recovered tRNA was suitable as a substrate for MetRS charging. The charging reaction (2 mL) contained approximately 200 nmoles of total tRNA and 10 nmoles of MetRS, along with 1 mM methionine, 2 mM ATP, 10 mM MgCl₂, 20 mM KCl, 4 mM DTT, 50 μg/mL BSA, and 50 mM Tris-HCl (pH 7.5). After incubating at 37°C for 10 minutes, a 4 μL sample was taken to measure charging efficiency as described by Gamper et al. ^54^. The remaining reaction mixture was further incubated at 37°C for 10 minutes with 1.7 μmoles of 10-formyltetrahydrofolate and 20 nmoles of methionyl formyl transferase to convert Met-tRNA to fMet-tRNA^fMet^. The reaction was stopped by adding a 0.1 volume of 2.5 M NaOAc (pH 5.0). After an extraction with an equal volume of phenol-chloroform-isoamyl alcohol (80:17:3, pH 5), the tRNA was precipitated with ethanol, resuspended in 300 μL of 25 mM NaOAc (pH 5.0), and stored at - 70°C. Around 200 nmoles of tRNA were recovered, with 45% being fMet-tRNA^fMet^.

### *In vitro* reconstitution of the 70S-fMet-tRNA^fMet^-mRNA complex

Purified *Mtb* C- 70S ribosomes (200 nM) were mixed with 1 µM each of fMet-tRNA^fMet^ and synthetic mRNA template (GGCAAGGAGGUAAAAAUGUUCAAAAAA) (IDT, USA) in polymix buffer (95 mM KCl, 5 mM NH4Cl, 5 mM Mg(OAc)2, 0.5 mM CaCl2, 8 mM putrescine, 1 mM spermidine, 5 mM potassium phosphate (KP) (pH 7.5) and 1 mM DTE)^55^. The reaction mixture was then incubated at 37°C for 30 minutes.

### Cryo-electron microscopy and image processing

Quantifoil R1.2/1.3 copper grids coated with holey carbon and an additional 2 nm carbon layer were further reinforced with a continuous carbon layer approximately 50 nm in thickness. These grids were subjected to a 30-second glow discharge in a plasma sterilizer. For sample vitrification, a Vitrobot IV (FEI) system was used. A 4 µL aliquot of the ribosome or ribosome-ligand complex was applied to the grids and allowed to incubate at 4°C with 100% humidity for 15 seconds, followed by a 4- second blotting step to remove excess sample. The grids were then rapidly plunge-frozen in liquid ethane. Intermediate resolution density maps reconstructed using cryo-EM images collected on an in-house JEOL3200FSC 300 KV microscope were used to confirm grid quality and sample composition. The cryo-EM images used for the final high-resolution reconstructions were captured using a Titan Krios electron microscope operating at 300 keV, equipped with a K3 direct electron detector (Gatan). The datasets were processed using cryoSPARC v3. Software ^56^. The details of data collection, processing, and model refinement for individual datasets are outlined below:

### For the Mtb C- 70S ribosomes

Micrographs were acquired at a nominal magnification of 64,000x, corresponding to a pixel size of 1.07 Å, with a total electron dose of 50.75 e-/Å². The defocus range for these images varied from −0.8 µm to −2.5 µm. A total of 13,499 movies were processed with patch-motion correction, and CTF parameters were estimated. Poor-quality micrographs were removed after manual curation, leaving 13,012 micrographs. From these, 5,471,811 particles were selected using template-based particle picking, with the template generated from a low-pass filtered map of the Mtb 70S ribosome ^57^. Subsequent rounds of 2D classification eliminated non-ribosomal particles (see fig. S3), retaining 3,460,467 particles for further processing via ab initio classification. This step resulted in three distinct particle classes, one representing 70S ribosomes (2,168,613 particles), one representing 50S subunits (761,965 particles), and the remainder classified as junk particles.

The 70S particles were further classified into five groups (vacant 70S, 70S-E-tRNA, 70S-P-tRNA, 70S-A-P-tRNA, and 70S-Mpy complexes) using a mask that focused on the interface between the large and small ribosomal subunits, encompassing the A, P, and E tRNA binding sites and the factor binding site. These were further classified using a mask around the NPET, identifying RTOF-bound and unbound states of vacant 70S, 70S-E-tRNA, and 70S-Mpy complexes. The RTOF-bound states were refined to resolutions of 2.8 Å, 2.7 Å, and 2.7 Å, respectively. No RTOF-bound states were observed for the 70S-P-tRNA and 70S- A-P-tRNA classes.

The 50S subunit class revealed a density in the NPET when visualized at lower thresholds. Masked 3D classification, focused on the NPET, generated five subclasses, with two showing clear RTOF density. The best-resolved RTOF class was refined to a 2.4 Å resolution.

### For the Mtb C- 70S-fMet-tRNA^fMet^-mRNA complex

Micrographs were collected at 105,000x magnification, corresponding to a pixel size of 0.84 Å, with a total dose of 51.65 e-/Å² and a defocus range of −0.5 to −2.5 µm. A total of 19,339 movies were aligned using patch-motion correction, followed by CTF parameter estimation. After manual curation, 19,104 micrographs were used for template-based particle picking, yielding 5,378,072 particles. After manual curation and several rounds of 2D classification to eliminate non-ribosomal particles, 2,809,935 particles were retained for ab initio classification into three classes. The 70S class comprised 705,168 particles, the 50S class contained 616,340 particles, and the remaining 1,488,427 particles were classified as either ribosomes with preferred orientations or junk (see fig. S6).

The 70S class was further processed using a mask around the LSU-SSU interface (encompassing the A, P, and E tRNA binding sites and the factor binding site). This resulted in two major classes: 70S-P-E-tRNA complexes (651,706 particles) and a smaller class of 70S-E-tRNA-RTOF complexes (53,462 particles). The 70S-P-E-tRNA class was further refined using a mask around the NPET, yielding RTOF-bound and unbound 70S-P-E-tRNA states, with the highest RTOF density map being refined to a final resolution of 3.3 Å. Finally, the 50S class was classified using a mask around the NPET, yielding RTOF-bound (304,680 particles) and unbound (311,660 particles) classes.

### Model building

The coordinates from the previously reported Mycobacterium tuberculosis ribosome structure (PDB: 7MT7, 70S) were fitted into the respective maps as rigid bodies using Chimera version 1.14 ^58^. Homology models for the C- ribosomal proteins S14, S18, L28 (Rv2058c and Rv0105c), and L33 were constructed with Swiss-model ^59^, utilizing the corresponding C- proteins from *M. smegmatis* (PDB: 6DZI) as templates ^8^. These models were docked into the relevant densities within the map using Chimera version 1.14 ^58^. Adjustments were made to the models based on cryo-EM density for better fitting using COOT ^60^, followed by refinement in PHENIX version 1.14 ^61^. The RTOF structure was identified and fitted as outlined in the Results section. Further refinements of the models were carried out in PHENIX version 1.14 ^61^. RNA and protein geometries were validated using MolProbity. Comprehensive statistics for EM reconstruction and model refinement are provided in tables S1–S2. The structural figures included in the manuscript were created using ChimeraX version 1.0 48 ^62–64^ and Chimera version 1.14 ^58^.

## Acknowledgements

This work was supported by grant to AKO and RKA (NIH: R01 AI132422). RKA also acknowledges support to his lab through NIH R01 grant Al155473. Authors thank Dr. Manjuli Sharma for discussion. Authors acknowledge Wadsworth Center’s and New York Structural Biology Center’s (NYSBC’s) 3D-EM facilities. Wadsworth Center is a contributing member of NYSBC, whose EM facility is supported by the Simons Foundation (SF349247).

## Author Contributions

YL purified Mtb ribosomes. HBG and Y-MH provided purified fMet-tRNA_i_^fMet^. SM and RKA designed structural experiments. SM prepared Mtb ribosome complexes with mRNA and tRNAs, performed detailed image processing of all complexes, discovered RTOF and identified its amino-acid sequence directly from cryo-EM structures by molecular modeling, identified gene encoding RTOF, and performed molecular analyses of all structures. SRM prepared and screened the cryo-EM grids, collected cryo-EM data on inhouse microscope, and obtained initial reconstructions. NKB helped in the in-house cryo-EM data collection and contributed to RTOF sequence analysis. NKB and RKA contributed to structural analyses and interpretation. SM and RKA wrote the manuscript, with help from NKB and AKO. All authors read, edited, and approved the manuscript.

## Data Deposition

Accession codes for the cryo-EM density maps and their corresponding atomic coordinates for the following four complexes that have been deposited in the Electron Microscopy Data Bank [https://www.ebi.ac.uk/pdbe/entry/emdb/] and the Protein Data Bank [https://www.rcsb.org/structure/], respectively, are EMD-47419 and PDB ID-9E1S for the 50S-RTOF complex, EMD-47384 and PDB ID-9E16 for the 70S-E-tRNA-RTOF complex, EMD-47876 and PDB ID-9EBB for the 70S-Mpy-RTOF complex, and EMD-47899 and PDB ID-9EC3 for the 70S-P-E-tRNAs-RTOF complex. Accession codes for the cryo-EM maps of two complexes that were obtained as control for qualitative comparison are also deposited in the Electron Microscopy Data Bank [https://www.ebi.ac.uk/pdbe/entry/emdb/] are EMD-47909 for the 70S-RTOF complex, and EMD-47910 for the 70S-P-E-tRNAs complex.

## Supplemental Materials

### Supplementary Figures

**Supplementary Figure 1.**
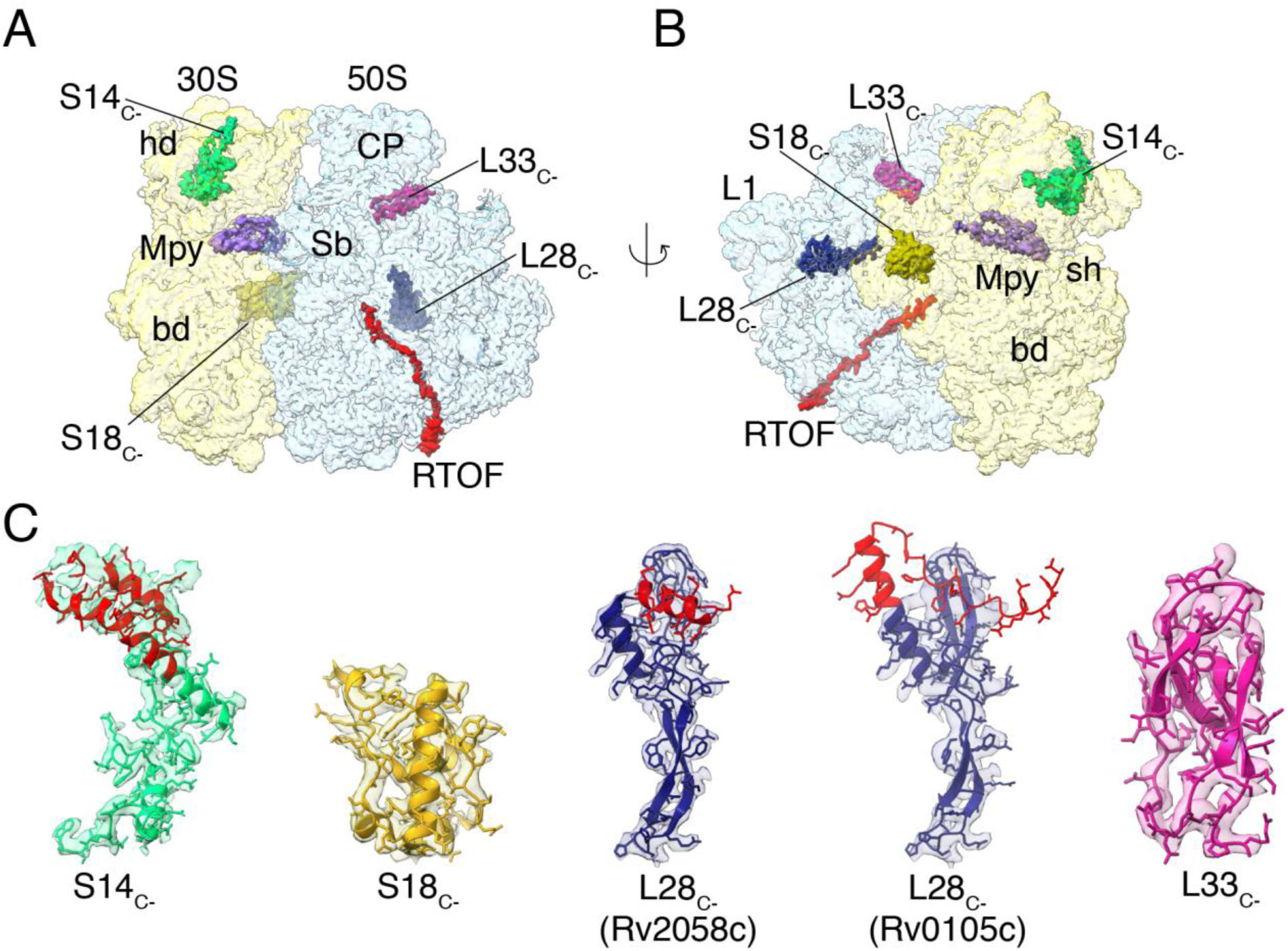
A 2.7 Å-resolution cryo-EM structure of the Mtb C− 70S ribosome bound to Mpy and RTOF proteins. (A, B) The C- 70S Mtb ribosome shown from two viewing directions: (A) 30S subunit (semitransparent yellow) and 50S subunit (semitransparent blue) in a side-by-side view, and (B) solvent-side view of the 30S subunit. Densities corresponding to the C− proteins—S14_C-_ (green), S18_C-_ (gold), L28_C-_ (slate blue), L33_C-_ (magenta)—as well as Mpy (violet) and RTOF (red) are highlighted in solid colors. (C) Modeling of all four C− ribosomal proteins into their respective cryo-EM densities. Extensions and insertions unique to the C− ribosomal proteins S14_C-_ and L28_C-_ are highlighted in red. A comparison of the fit between both paralogs of L28_C-_ indicates that the density corresponds to Rv2058c.

**Supplementary Figure 2.**
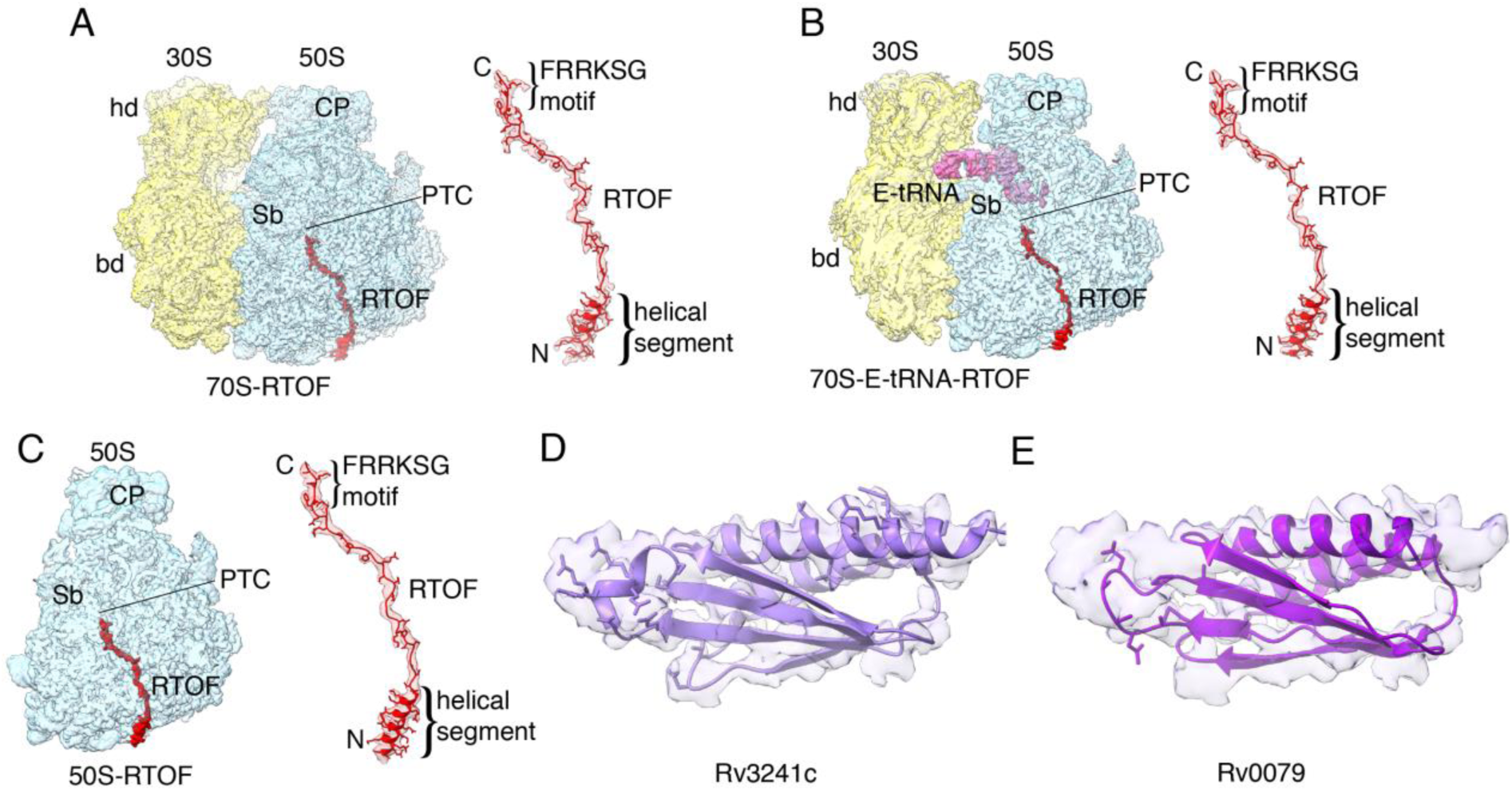
Structures of RTOF-bound ribosomal states determined in this study. (A) A 2.8 Å cryo-EM structure of the 70S-RTOF complex. (B) A 2.7 Å cryo-EM structure of the 70S-E-tRNA-RTOF complex. (C) A 2.4 Å cryo-EM structure of the 50S-RTOF complex. The corresponding RTOF densities in each structure are displayed on the right-hand side of each panel. (D, E) A comparison of the fit between the two Mpy homologs suggests that the density corresponds to the N-terminal domain of Rv3241c.

**Supplementary Figure 3.**
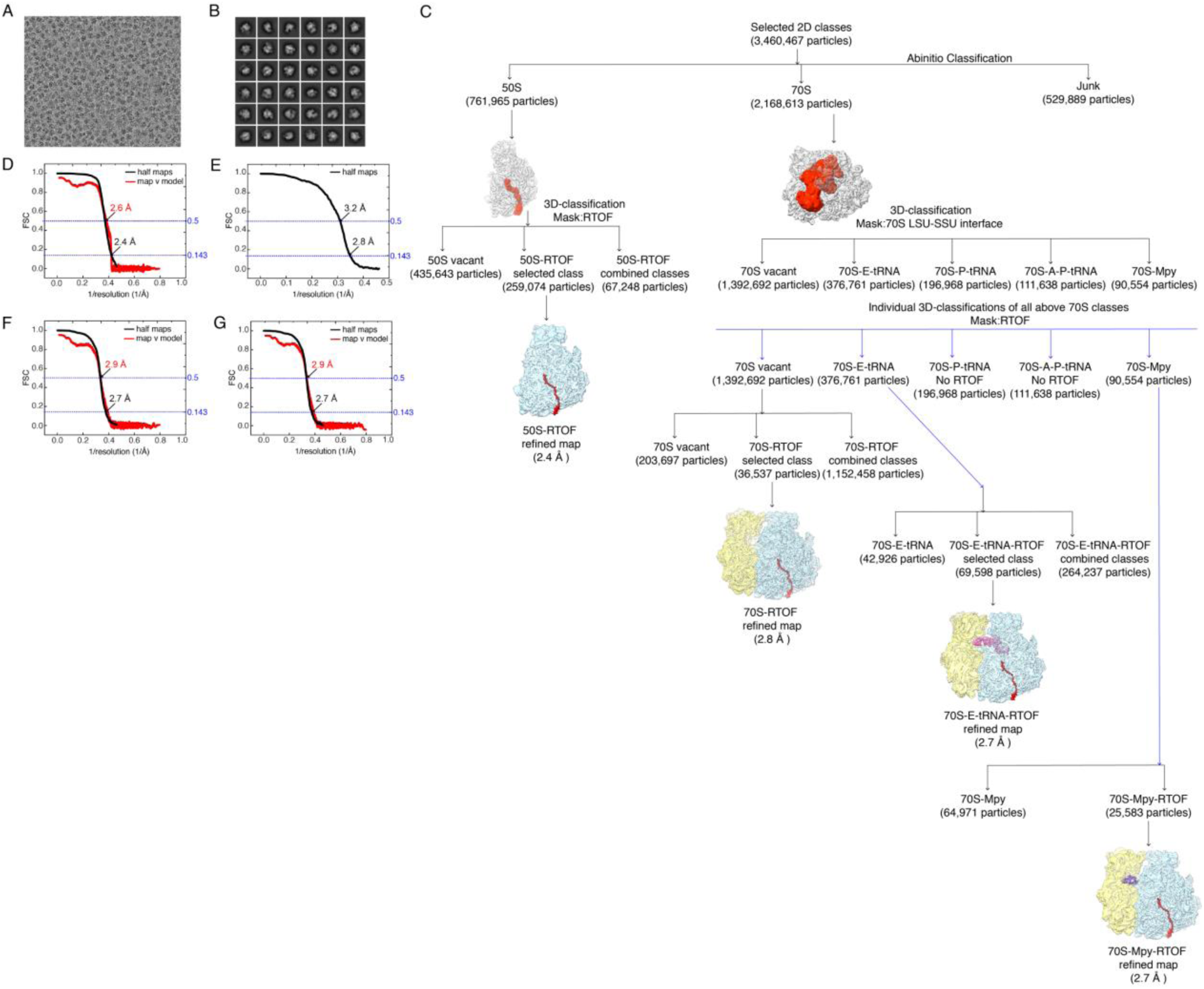
Image processing of Mtb C− 70S ribosomes. (A) A representative micrograph from the cryo-EM dataset. (B) Representative 2D-class averages selected for further data processing. (C) Flowchart detailing the 3D classifications and refinements performed on the selected particles. The final four maps used for model building and interpretation are displayed. (D-G) Gold-standard Fourier shell correlation (FSC) curves (black) overlaid with map-to-model FSC curves (red) for the (D) 50S-RTOF, (E) 70S-RTOF, (F) 70S-E-tRNA-RTOF, and (G) 70S-Mpy-RTOF structures.

**Supplementary Figure 4.**
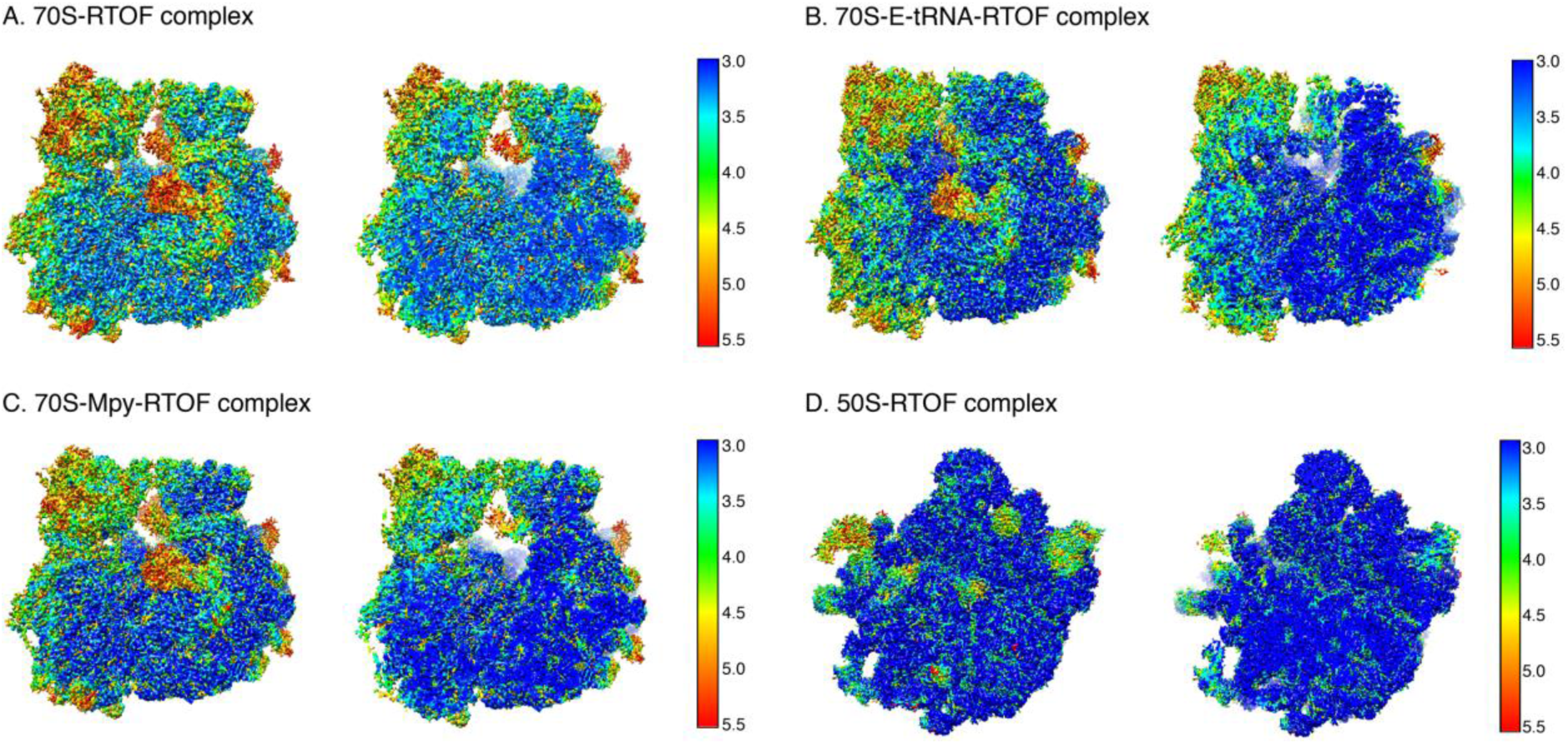
Local resolution of the cryo-EM maps obtained from Mtb C− 70S ribosomes. Local resolution of the maps was estimated using ResMap ^1^. The densities are color-coded according to the local resolution, with the corresponding color keys provided alongside. The left-hand-side snapshot in each panel (A-D) represents the complete map, while the right-hand-side snapshot shows a slice through center of the corresponding map.

**Supplementary Figure 5.**
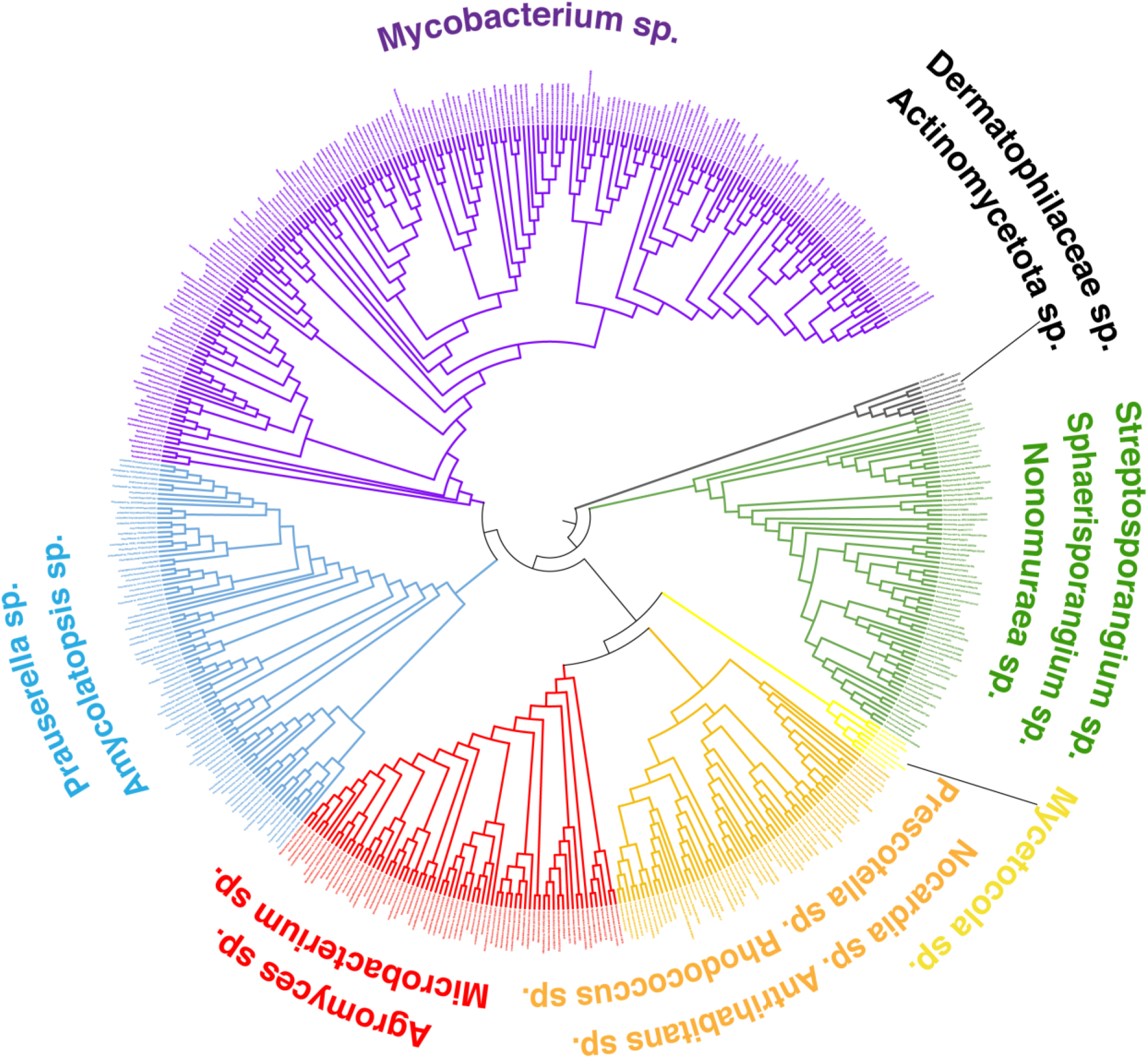
Phylogenetic tree illustrating sequence variation and evolutionary relationships among RTOF homologs within Actinobacteria. Using NCBI Delta-BLAST, 500 sequences most similar to Rv1155a were identified. These sequences were aligned with COBALT^2^, and a phylogenetic tree was constructed based on the alignment. The tree was visualized and annotated using the Interactive Tree of Life (iTOL)^3^ online tool.

**Supplementary Figure 6.**
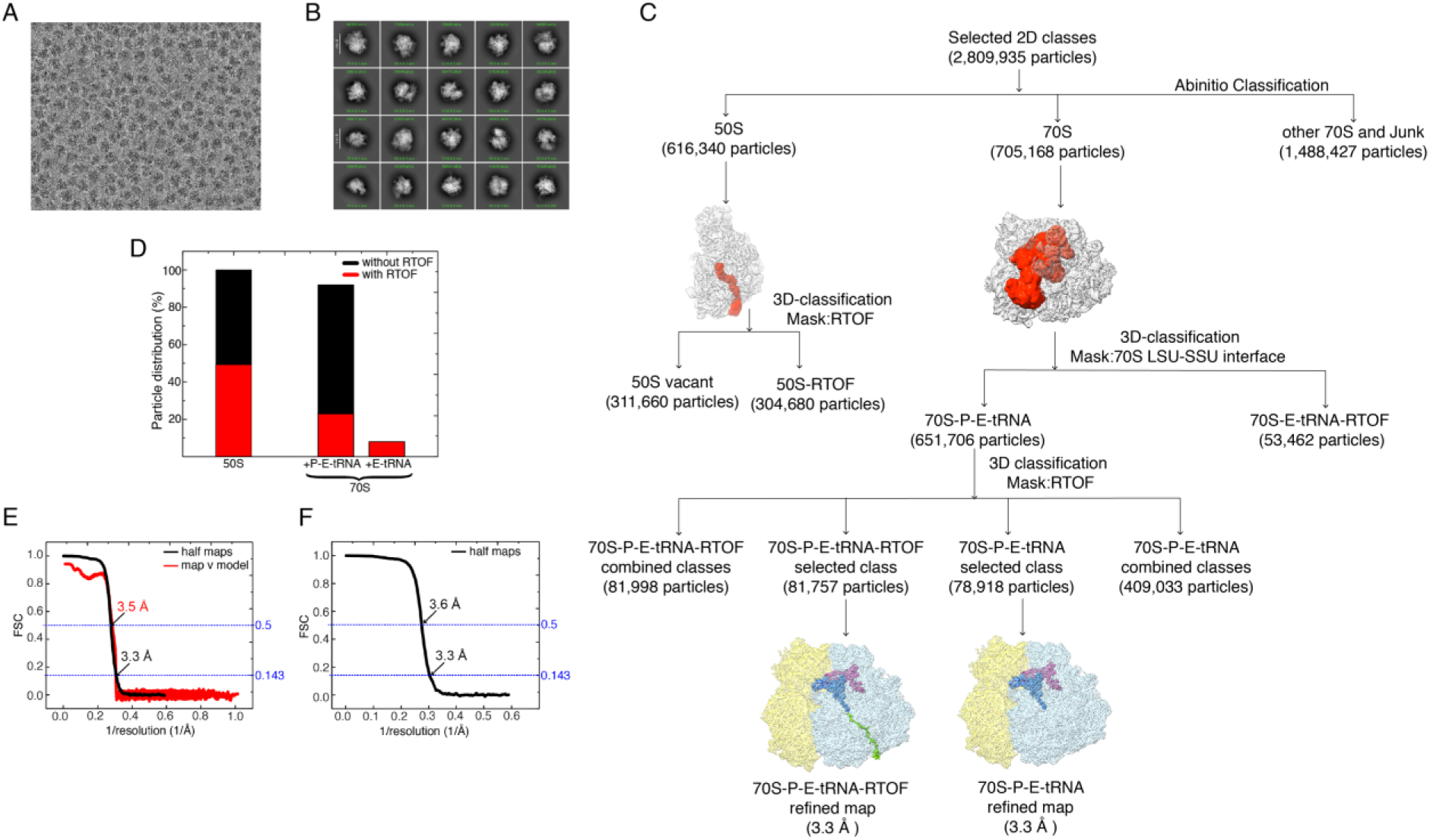
Image processing of the Mtb C− 70S-fMet-tRNA^fMet^-mRNA complex. (A) A representative micrograph from the cryo-EM dataset. (B) Representative 2D- class averages selected for further data processing. (C) Flowchart detailing the 3D classifications and refinements performed on the selected particles. The final two maps used for model building and interpretation are displayed. (D) Plot showing the percent distribution of RTOF-bound (red) and unbound (black), 50S, 70S-fMet-tRNA^fMet^-E-tRNA (+P-E-tRNA), and 70S-E-tRNA (+E-tRNA) complexes. (E, F) Gold-standard Fourier shell correlation (FSC) curves (black) overlaid with map-to-model FSC curves (red) for the (D) 70S-P-E-tRNA- RTOF and (E) 70S-P-E-tRNA structures.

**Supplementary Figure 7.**
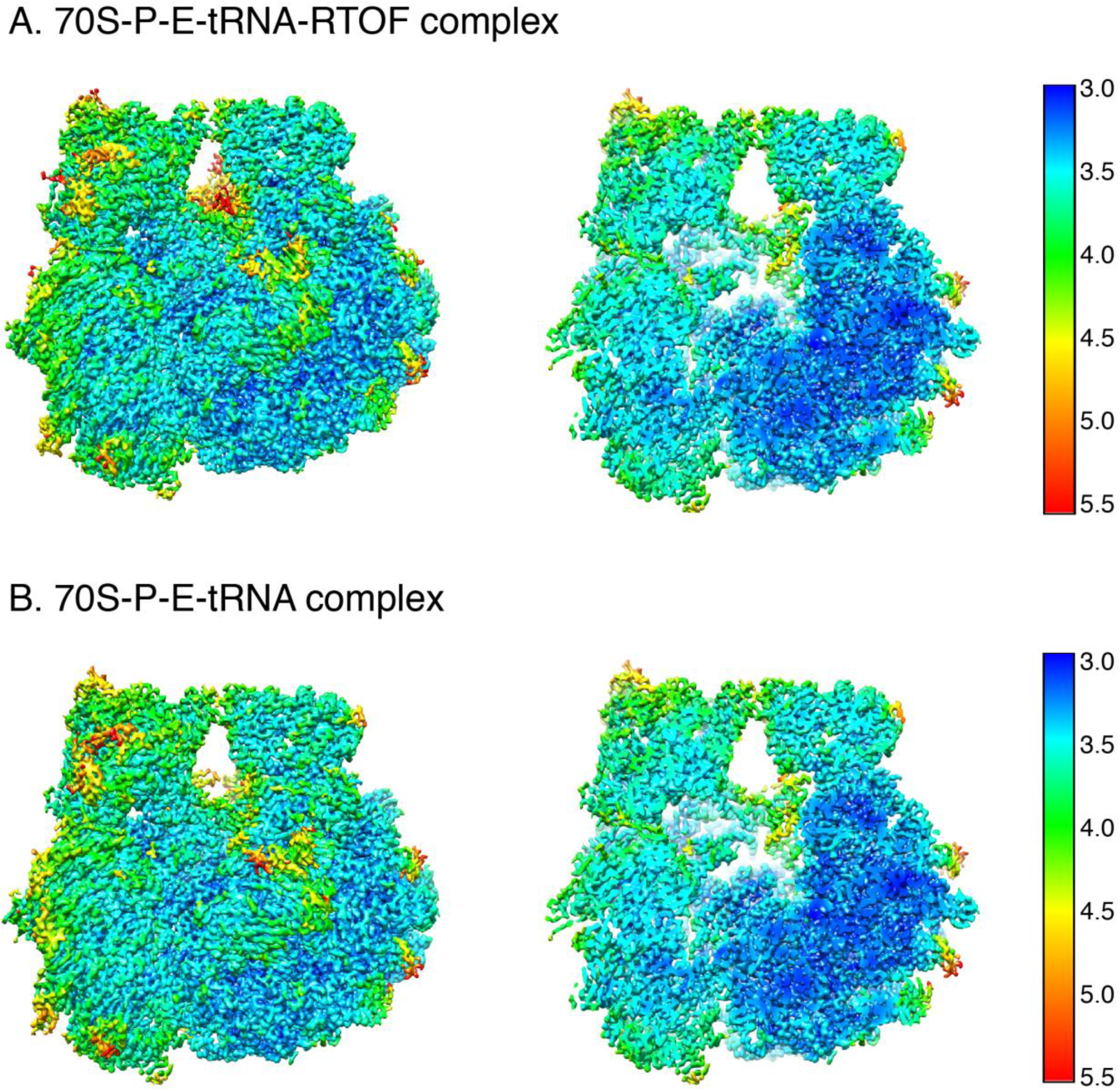
Local resolution of the cryo-EM maps obtained from the Mtb C− 70S-fMet-tRNA^fMet^-mRNA complex. Local resolution of the maps was estimated using ResMap ^1^. The densities are color-coded according to the local resolution, with the corresponding color keys provided alongside. The first left-hand-side snapshot in panels (A-B) represents the complete map, while the right-hand-side snapshot shows slices through the same map.

**Supplementary Figure 8.**
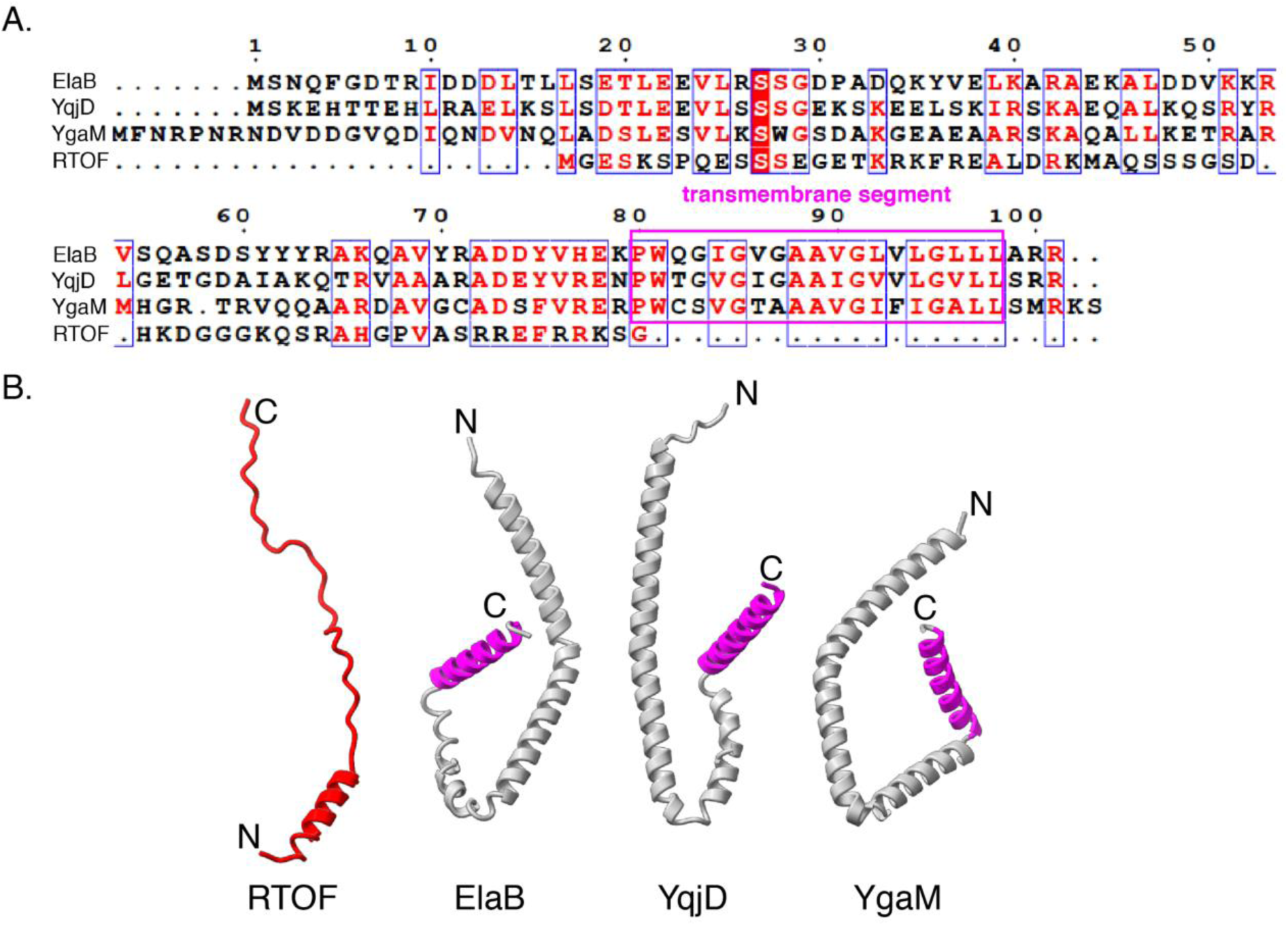
(A) Sequence alignment of putative ribosomal NPET-associated hibernation factors from *E. coli*—ElaB, YqjD, and YgaM—along with Mtb RTOF. Sequence similarity and identity are highlighted in red. (B) Comparison of the cryo-EM structure of RTOF, as determined in this study, with the AlphaFold models of ElaB, YqjD, and YgaM. The transmembrane segments of the latter proteins are indicated in magenta.

**Supplementary Figure 9.**
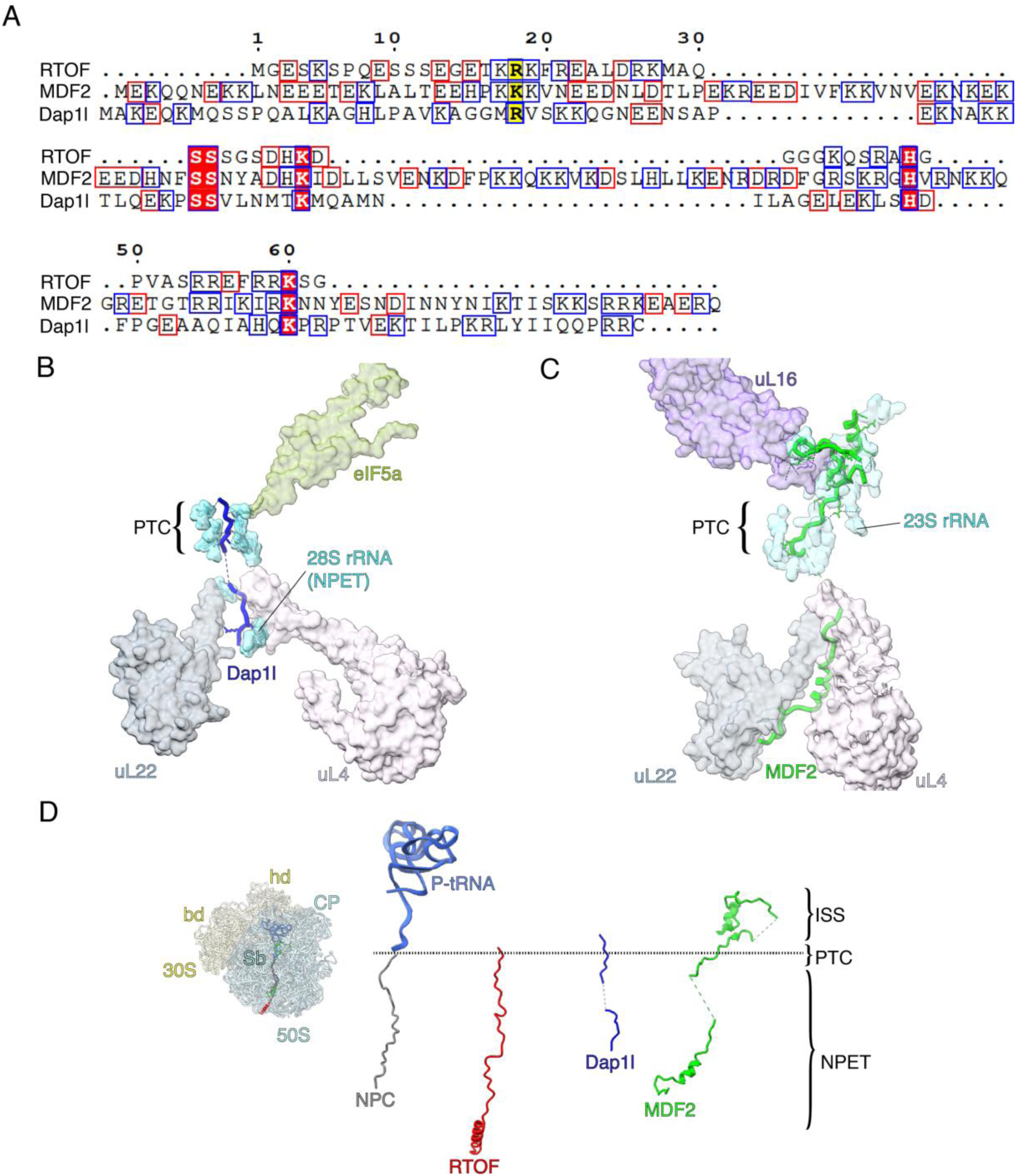
(A) Sequence alignment of Mtb RTOF with two known eukaryotic NPET-associated hibernation factors, Dap1l (*Xenopus*) and MDF2 (*Vairimorpha necatrix*). Sequence identity is highlighted with red-filled boxes. Charged amino acids—Glu (E) and Asp (D)—are outlined with red boxes, while Arg (R), Lys (K), and His (H) are outlined with blue boxes. (B, C) Overall interactions of Dap1l and MDF2 with NPET components, respectively ^4,5^. (D) Comparison of the extent to which the hibernation factors—RTOF (red), Dap1l (blue), and MDF2 (green)—extend into the inter-subunit space (ISS), as compared to a nascent polypeptide chain (NPC, grey).

**Supplementary Table 1.**
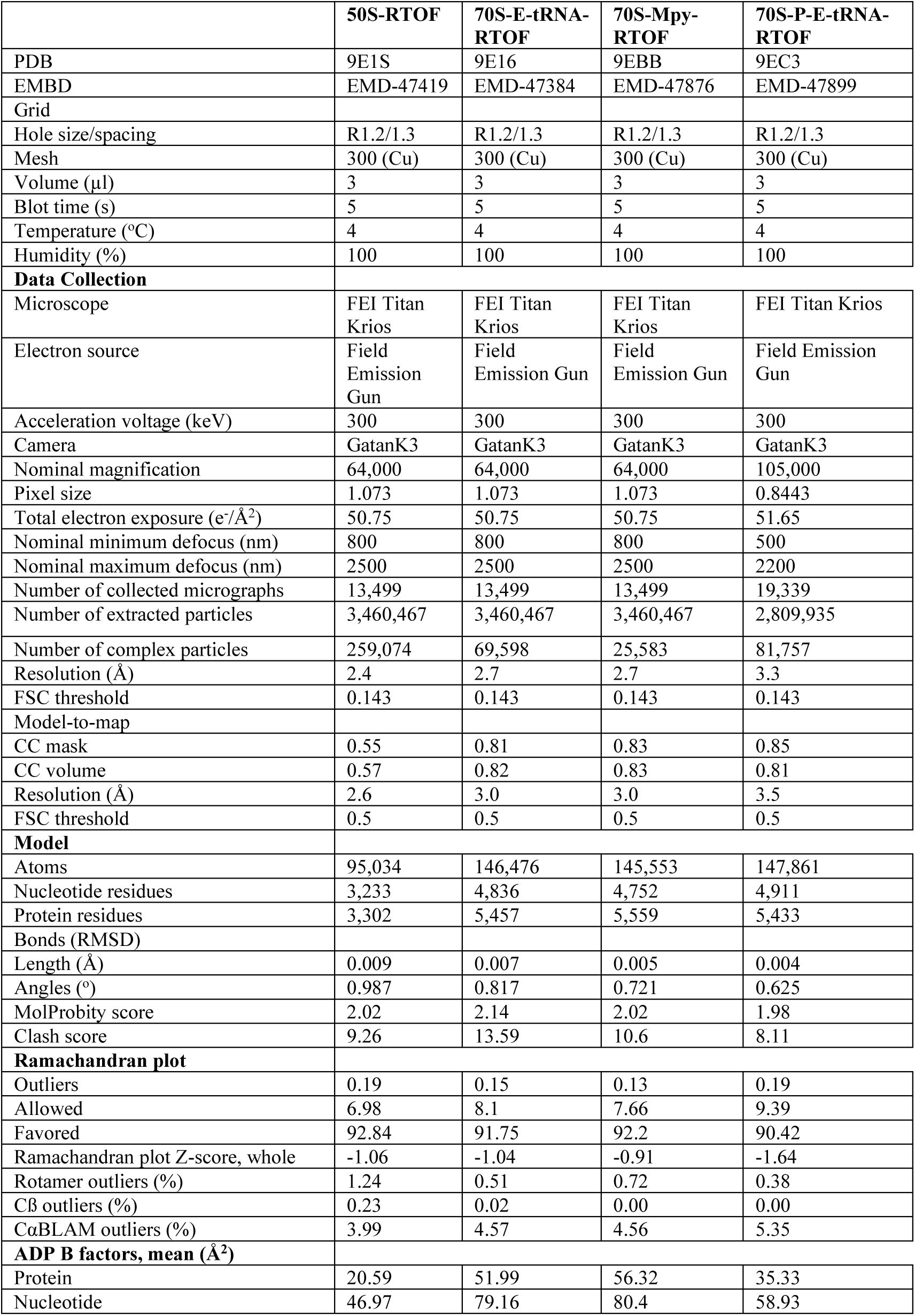
Grid preparation, data collection and refinement parameters for the ribosome-RTOF complexes.

**Supplementary Table 2.** Grid preparation, data collection and refinement parameters.

|  | **70S-RTOF** | **70S-P-E-tRNA** |
| --- | --- | --- |
| EMBD | EMD-47909 | EMD-47910 |
| Grid |  |  |
| Hole size/spacing | R1.2/1.3 | R1.2/1.3 |
| Mesh | 300 (Cu) | 300 (Cu) |
| Volume (μl) | 3 | 3 |
| Blot time (s) | 5 | 5 |
| Temperature (°C) | 4 | 4 |
| Humidity (%) | 100 | 100 |
| **Data Collection** |  |  |
| Microscope | FEI Titan Krios | FEI Titan Krios |
| Electron source | Field Emission Gun | Field Emission Gun |
| Acceleration voltage (keV) | 300 | 300 |
| Camera | GatanK3 | GatanK3 |
| Nominal magnification | 64,000 | 105,000 |
| Pixel size | 1.073 | 0.8443 |
| Total electron exposure (e<sup>-</sup>/Å<sup>2</sup>) | 50.75 | 51.65 |
| Nominal minimum defocus (nm) | 800 | 500 |
| Nominal maximum defocus (nm) | 2500 | 2200 |
| Number of collected micrographs | 13,499 | 19,339 |
| Number of extracted particles | 3,460,467 | 2,809,935 |
| Number of complex particles | 36,537 | 78,918 |
| Resolution (Å) | 2.8 | 3.3 |
| FSC threshold | 0.143 | 0.143 |

